# Mutually suppressive roles of KMT2A and KDM5C in behaviour, neuronal structure, and histone H3K4 methylation

**DOI:** 10.1101/567917

**Authors:** Christina N. Vallianatos, Brynne Raines, Robert S. Porter, Katherine M. Bonefas, Michael C. Wu, Patricia M. Garay, Katie M. Collette, Young Ah Seo, Yali Dou, Catherine E. Keegan, Natalie C. Tronson, Shigeki Iwase

## Abstract

Histone H3 lysine 4 methylation (H3K4me) is extensively regulated by numerous writer and eraser enzymes in mammals. Nine H3K4me enzymes are associated with neurodevelopmental disorders to date, indicating their important roles in the brain. However, interplay among H3K4me enzymes during brain development remains largely unknown. Here, we show functional interactions of a writer-eraser duo, *KMT2A* and *KDM5C*, which are responsible for Wiedemann-Steiner Syndrome (WDSTS), and mental retardation X-linked syndromic Claes-Jensen type (MRXSCJ), respectively. Despite opposite enzymatic activities, the two mouse models deficient for either *Kmt2a* or *Kdm5c* shared reduced dendritic spines and increased aggression. Double mutation of *Kmt2a* and *Kdm5c* clearly reversed dendritic morphology, key behavioral traits including aggression, and partially corrected altered transcriptomes and H3K4me landscapes. Thus, our study uncovers common yet mutually suppressive aspects of the WDSTS and MRXSCJ models and provides a proof of principle for balancing a single writer-eraser pair to ameliorate their associated disorders.

## Introduction

Dysregulation of histone methylation has emerged as a major contributor to neurodevelopmental disorders (NDDs) such as autism spectrum disorders and intellectual disabilities (1). Histone methylation can be placed on a subset of lysines and arginines by histone methyltransferases (writer enzymes) and serves as a signaling platform for a variety of nuclear events including transcription (2). Reader proteins specifically recognize methylated histones, thereby converting methylation signals into higher-order chromatin structures. Histone methylation can be removed by a set of histone demethylases (eraser enzymes). Pathogenic variants in all three classes of methyl-histone regulators cause NDDs, indicating critical, yet poorly understood roles of histone methylation dynamics in brain development and function (3).

Histone H3 lysine 4 methylation (H3K4me) is one of the most well-characterized histone modifications. H3K4me is primarily found at transcriptionally active areas of the genome. The three states, mono-, di-, and tri-methylation (H3K4me1-3), uniquely mark gene regulatory elements and play pivotal roles in distinct steps of transcription. While H3K4me3/2 are enriched at transcriptionally engaged promoters, H3K4me1 is a hallmark of transcriptional enhancers (4, 5). At promoters, H3K4me3 contributes to recruitment of general transcription machinery TFIID and RNA polymerase II (6, 7). H3K4me1 at enhancers can be recognized by BAF, an ATP-dependent chromatin remodeling complex (8).

H3K4me is extensively regulated by more than seven methyltransferases and six demethylases in mammals (9). Consistent with the important roles of H3K4me in transcriptional regulation, genomic distribution of H3K4me appears highly dynamic during brain development (10). However, the contributions of each of the 13 enzymes in the dynamic H3K4me landscapes of the developing brain remain largely unknown. Strikingly, genetic alterations in nine H3K4me enzymes and at least two H3K4me readers have been associated with human NDDs to date, indicating the critical roles of H3K4me balance (10) (Figure 1A). These human conditions can be collectively referred to as brain H3K4 methylopathies and point to non-redundant yet poorly understood roles of these genes controlling this single post-translational modification for faithful brain development. Of note, some of these enzymes can have non-enzymatic scaffolding function (11) as well as non-histone substrate (12); therefore, these disorders may potentially involve mechanism outside histone H3K4 methylation.

**Figure 1.**
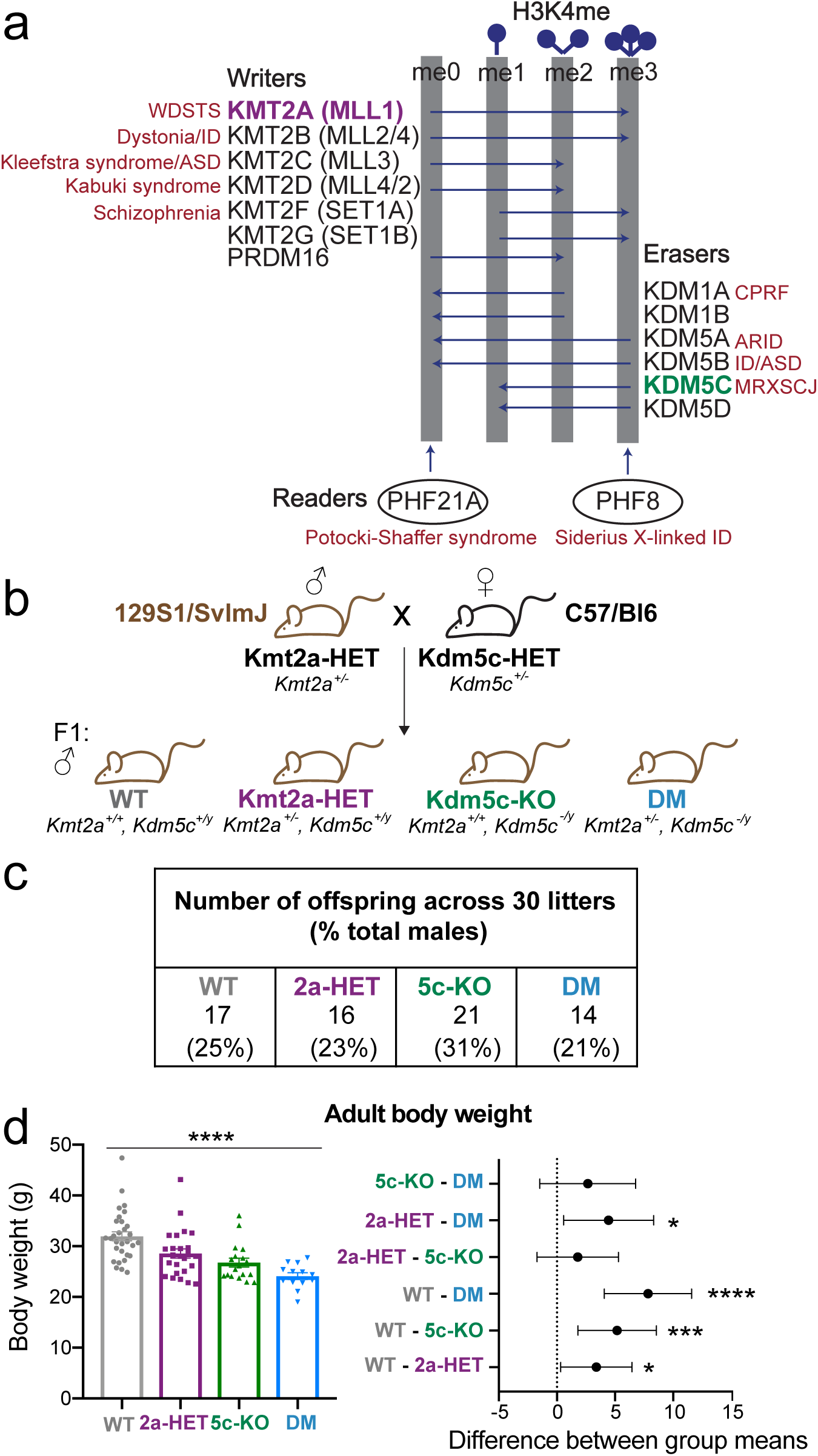
The H3K4 methylopathies and generation of the *Kmt2a-Kdm5c* double-mutant (DM) mouse. (A) Histone H3 lysine 4 (H3K4me) methyltransferases (writers) and demethylases (erasers) depicted by their ability to place or remove H3K4me. Reader proteins recognizing specific H3K4me substrates (arrows) are depicted below. Genes are listed next to their associated neurodevelopmental disorder. KMT2A and KDM5C are highlighted in purple and green, respectively. WDSTS: Weideman-Steiner Syndrome; ID: intellectual disability; ASD: autism spectrum disorder, CPRF: cleft palate, psychomotor retardation, and distinctive facial features; ARID: autosomal recessive ID; MRXSCJ: mental retardation, X-linked, syndromic, Claes-Jensen type. **(B)** Mouse breeding scheme crossing congenic 129S1/SvlmJ *Kmt2a*-heterozygous males with congenic C57/Bl6 *Kdm5c*-heterozygous females, resulting in F1 generation mice. Only males were used in this study. **(C)** Numbers of male offspring across 30 litters, showing Mendelian ratios of expected genotypes. **(D)** Left panel: Body weight of adult mice > 2 months of age (mean ± SEM, \*\*\*\**p* < 0.0001 in one-way ANOVA). Right panel: Difference between group means of weight (mean ± 95% confidence intervals, \**p*<0.05, \*\**p*<0.01, \*\*\**p*<0.001, \*\*\*\**p*<0.0001 in Tukey’s multiple comparison test).

As histone modifications are reversible, one can, in theory, correct an imbalance by modulating the writers or erasers. Chemical inhibitors of histone deacetylases (HDACs) have been successfully used to rescue phenotypes in mouse models of NDDs. HDAC inhibitors were able to ameliorate learning disabilities in mouse models of Rubinstein-Taybi and Kabuki syndromes, which are deficient for CREBBP or KMT2D, writer enzymes for histone acetylation or H3K4me, respectively (13, 14). However, the HDAC inhibitors, such as SAHA and AR-42, used in these studies interfere with multiple HDACs (15), which could potentially result in widespread side effects. Given the non-redundant roles of the H3K4me enzymes, a more specific perturbation is desirable.

In order to achieve specific modulation of H3K4me, an important first step is to delineate functional relationships between the H3K4 enzymes. Focus of the work is on a pair of NDD-associated writer/eraser enzymes: KMT2A and KDM5C. Haploinsufficiency of *KMT2A* underlies Weidemann-Steiner Syndrome (WDSTS), characterized by developmental delay, intellectual disability, characteristic facial features, short stature, and hypotonia (16). Loss of KDM5C function defines Mental Retardation, X-linked, syndromic, Claes Jensen type (MRXSCJ), in which individuals display an intellectual disability syndrome with aggression, short stature, and occasional autism comorbidity (17). Mouse models have provided experimental support for causative impacts of KMT2A and KDM5C deficiencies in impaired cognitive development (18–22). Social behavior and neuronal structure in *Kmt2a*-KO mice however have not been characterized.

In the present work, we tested whether modulating a single H3K4me writer or eraser can ameliorate the neurodevelopmental symptoms observed in the WDSTS and MRXSCJ mouse models. We generated *Kmt2a*-, *Kdm5c*-double mutant (DM) mice, and performed systematic comparisons between wild-type (WT), single mutants, and DM mice.

## Results

### KMT2A and KDM5C co-exist broadly in the brain

We first examined expression patterns of KMT2A and KDM5C using publicly available resources, and found the two genes are broadly expressed throughout brain regions of adult mice and humans (Supplementary Figure 1). *Kmt2a* and *Kdm5c* are expressed at comparable levels in all major excitatory and inhibitory neuron subtypes as well as glia cells in mouse visual cortices (Supplementary Figure 1A), and also throughout mouse brains (Supplementary Figure 1B). Consistently, developing and aging human brains express *KMT2A* and *KDM5C* at high, steady levels (Supplementary Figure 1C). Thus, both writer and eraser are co-expressed across brain cell types, regions, and developmental stages in both humans and mice.

### Generation of *Kmt2a*-*Kdm5c* double-mutant (DM) mice

To test genetic interaction of *Kmt2a* and *Kdm5c*, we generated *Kmt2a*-*Kdm5c* double-mutant (DM) mice. Experimental mice were F1 hybrids of the two fully congenic strains: 129S1/SvImJ *Kmt2a*^+/-^ males (23) and C57BL/6J *Kdm5c*^+/-^ females (21) (Figure 1B). This cross resulted in the following genotypes of male mice in an identical genetic background: wildtype (WT); *Kmt2a* heterozygote (*Kmt2a*-HET: *Kmt2a*^+/-^), *Kdm5c* hemizygous knock-out (*Kdm5c*-KO: *Kdm5c*^-/y^), and *Kmt2a*-*Kdm5c* double-mutant (DM: *Kmt2a*^+/-^, *Kdm5c*^-/y^), thereby allowing us to perform a comparison between the WDSTS model (*Kmt2a*-HET), the MRXSCJ model (*Kdm5c*-KO), and their composite (DM). We focus on males, because MRXSCJ predominantly affects males and *Kdm5c*-heterozygous female mice exhibit only minor cognitive deficits (22).

These mice were born at expected Mendelian ratios of 25% per genotype, demonstrating the DM mice were not synthetic lethal (Figure 1C). Genotypes were confirmed at RNA and DNA levels (Supplementary Figure 1A-C), and protein level for KDM5C (Supplementary Figure 1D). Brain anatomy showed no gross deformities in any of the genotypes (Supplementary Figure 1E). Both *Kmt2a*-Het and *Kdm5c*-KO mice showed significant body weight reduction compared to WT (Figure 1D and Supplementary Figure 1F, One Way ANOVA: F(3, 55) = 10.28, *p* < 1.0 x 10^-4^, Tukey’s multiple comparison test: WT vs 2A: *p* = 0.008, WT vs 5C: *p* = 0.008). DM body weight was significantly smaller compared to WT (DM vs WT: *p* < 1.0 x 10^-4^). Thus, loss of *Kdm5c* and *Kmt2a* heterozygosity both led to growth retardation, which was not corrected but rather slightly exacerbated in DM mice. Note that for all the four-way comparisons in this study, we first represent the *p*-values from one-way ANOVA tests of genotype-phenotype interaction with histograms. We then report *p*-values from post-hoc tests of all six genotype comparisons with 95% confidence intervals of group mean differences.

### Memory impairments in *Kdm5c*-KO were ameliorated in DM

We first sought to determine the effect of loss of *Kmt2a* and/or *Kdm5c* on mouse behavior through a battery of behavioral tests. Learning and memory was measured by two independent tests, contextual fear conditioning (CFC) and novel object recognition (NOR). In CFC, we observed a significant effect of genotype (CFC: *F* (3,60) = 4.133, *p* = 0.010). In accordance with previous findings (21, 22), *Kdm5c*-KO showed significant deficits in associative fear memory in CFC (Figure 2A, WT vs 5C: *p* = 0.017, 2A vs 5C: *p* = 0.0.024). Previous work reported that homozygous deletion of *Kmt2a* in excitatory hippocampal neurons leads to impaired fear memory in the CFC (20). In contrast, *Kmt2a*-HET mice showed no deficits in either CFC or NOR (Figure 2A-B) (CFC: *p* = 1.000), indicating that *Kmt2a*-heterozygosity does not lead to learning impairment measured in these assays. Importantly, DM mice did not differ from WT mice (Figure 2A) (*p* = 0.923), suggesting that *Kmt2a* heterozygosity rescues CFC memory deficits of *Kdm5c*-KO mice.

**Figure 2.**
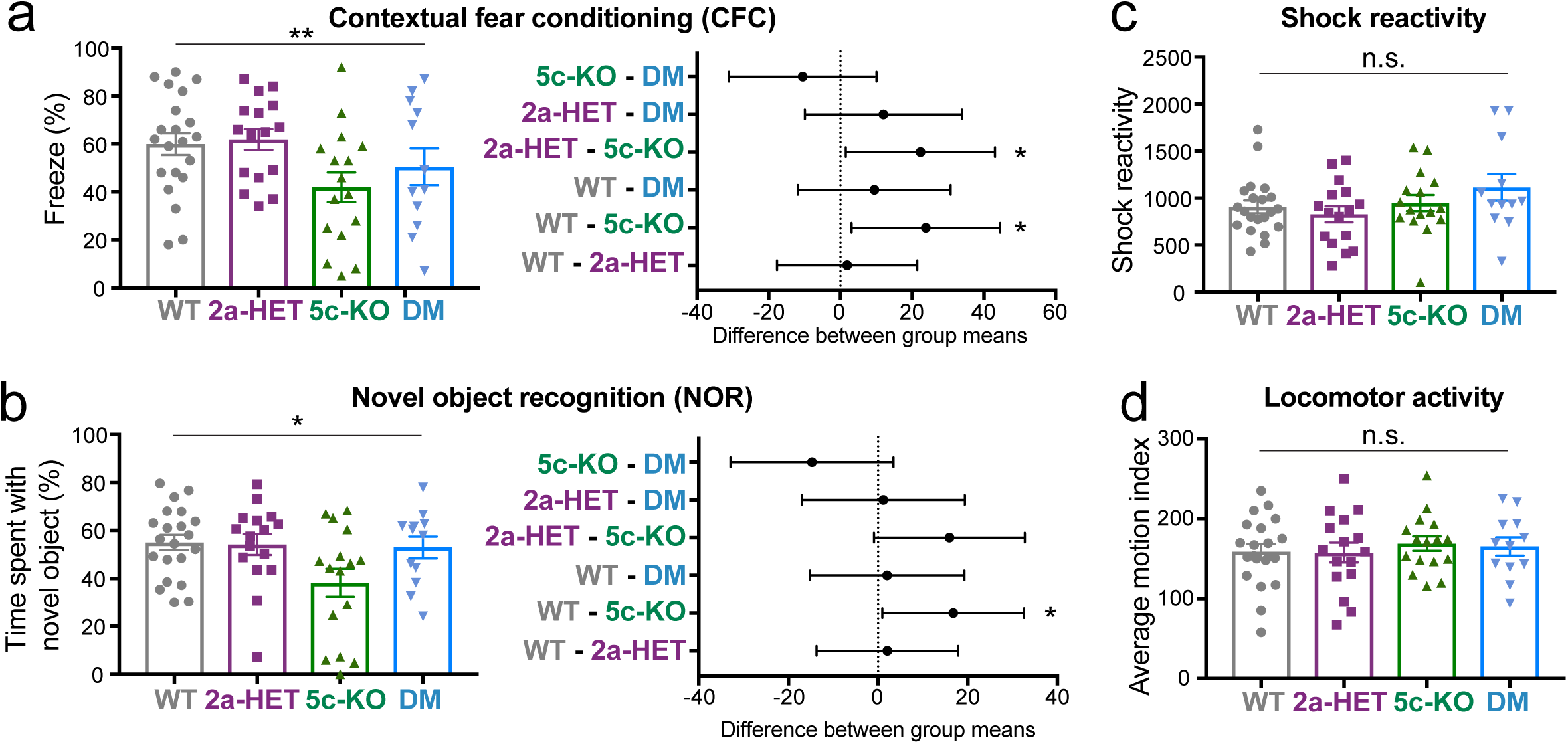
Deficit of memory-related behavior in *Kdm5c*-KO and its rescue in DM. (A) Contextual fear conditioning test. Left panel: Freezing levels after shock on test day (mean ± SEM, \**p* < 0.05). Right panel: Difference between group means of freezing (mean ± 95% confidence intervals, \**p* < 0.05). **(B)** Novel object recognition test. Left panel: Preference for novel versus familiar object (mean ± SEM, \**p* < 0.05). Right panel: Difference between group means of freeze response (mean ± 95% confidence intervals, \**p*<0.05). **(C)** Response to mild foot-shock (mean ± 95% confidence intervals, no statistical significance [n.s.],). **(D)** Locomotor activity (mean ± 95% confidence intervals, no statistical significance [n.s.]). N=21 WT, N=16 *Kmt2a*-HET, N=16 *Kdm5c*-KO, and N=12 DM animals were used for all studies.

A similar pattern emerged in NOR, where we observed a significant main effect of genotype (*F*(3,64) = 3.20, *p* = 0.030). Here, *Kdm5c-*KO mice showed significantly less preference for the novel object than other genotypes (Figure 2B) (WT *vs* 5C: *p =* 0.041; 2a *vs* 5C: p = 0.091). Consistent with our CFC results, neither *Kmt2a*-HET mice nor DM mice differed from WT mice (*p* = 1.000), suggesting that *Kmt2a* heterozygosity does not impair memory, but can rescue impairments of *Kdm5c*-KO. Importantly, WT, *Kmt2a*-Het, and DM mice showed preference for the novel object (*Z* = 2.029; *p* = 0.041). Nevertheless, as this was not a strong preference, it is likely that non-mnemonic effects, including anxiety processes, also contributed to avoidance-like behavior in *Kdm5c*-KO mice. Differences between genotypes in memory tasks were not attributable to locomotor activity or shock responsiveness, as none of these parameters showed significant differences among the genotypes (Figure 2C-D).

### Social behavior in the single and double mutants

We next examined social behavior using the three independent behavioral paradigms. First, social interaction was tested with the three-chambered preference test, with an overall effect of genotype (F(3,60) = 3.726, *p* = 0.016). WT mice showed a robust preference for the novel mouse over the toy mouse (*Z* = 2.97, *p* = 0.003). Consistent with previous studies (21), *Kdm5c*-KO mice exhibited significantly less preference for the stranger mouse compared with WT (WT vs 5C: *p=* 0.023); whereas *Kmt2a*-HET mice showed no differences from WT (Figure 3A, WT vs 2A: *p* = 0.563) (20). Similar to *Kdm5c*-KO, DM mice showed a strong trend towards less time with stranger mice compared to WT (WT vs DM: *p* = 0.077). No difference was detected between *Kdm5c*-KO and DM (5C vs DM: *p=* 1.000), indicating that *Kmt2a* heterozygosity does not alter social preference or rescue the deficit of *Kdm5c*-KO.

**Figure 3.**
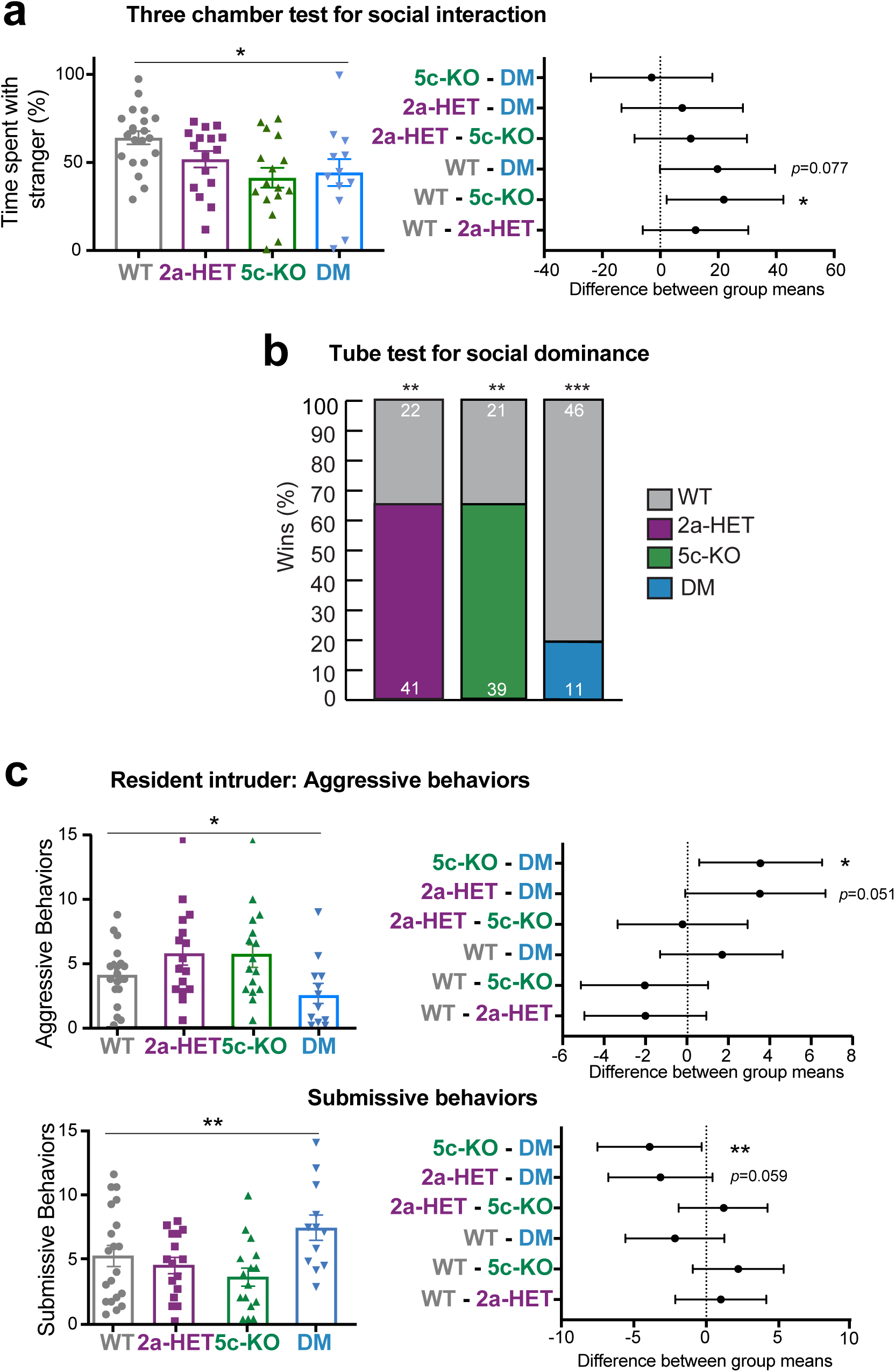
Differential impacts of double mutation in social behavior. (A) Three chamber test for social interaction. Left panel: preference for stranger versus toy mouse (mean ± SEM, \**p* < 0.05). Right panel: Difference between group means of preference (mean ± 95% confidence intervals, \**p*<0.05). **(B)** Tube test for social dominance. Proportion of wins in matches of each mutant versus WT. Numbers on colored bars represent total number of wins for WT (grey, above) or each mutant (below) in every matchup. \*\**p*<0.01, \*\*\**p*<0.001, Exact binomial test. **(C)** Resident intruder test. Left panel: average number of all aggressive and submissive behaviors (mean ± SEM, \**p* < 0.05, \*\**p* < 0.01). Right panel: Difference between group means of aggressive and submissive behaviors (mean ± 95% confidence intervals, \**p* < 0.05, \*\**p* < 0.01). N=21 WT, N=16 *Kmt2a*-HET, N=16 *Kdm5c*-KO, and N=12 DM animals were used for all studies.

In tests of social dominance (Figure 3B), both *Kmt2a*-HET and *Kdm5c*-KO mice won more frequently against WTs (2A vs WT: 60.9%, *p* = 0.091, 5C vs WT: 68.4%, *p* = 0.008). In contrast, DM animals lost more than 80% of their bouts against WT (DM vs WT: *p* = 1.47 x 10^-5^). Although DM mice were slightly smaller compared to single mutants (Figure 1D), this is unlikely to drive submissive behaviors, as body mass has been shown to have minimal impact on social hierarchy unless excess difference (> 30%) is present between animals (24), which is not the case in our study (Figure 1D) (2A vs WT:11%, 5C vs WT: 17%, DM vs WT: 25%). These results indicate that the two single mutants share heightened social dominance and the double mutation reverses the social dominance.

Lastly, in the resident-intruder test, similar to other behavioral paradigms, overall effects of genotype were evident in both aggressive and submissive behavior (Figure 3C, aggressive behavior: *F*(3,60) = 3.642, *p* = 0.018; aggression x genotype: *F*(12,240) = 1.853, *p* = 0.041; submissive behavior: (*F*(3,60) = 4.335, *p* = 0.008). The genotype effect on aggressive behaviors inversely correlated with that of submissive behavior, reinforcing the changes in specific behaviors rather than general locomotor activity. Both *Kmt2a*-HET and *Kdm5c*-KO showed a trend of increased aggression and decreased submission when we combined frequency of all aggressive or submissive behavior types compared to WT (Figure 3C).

DM mice showed significantly reduced overall aggression compared to the two single mutants (Fig. 3C, DM vs 5C: *p* = 0.045, DM vs 2A: *p* = 0.051). Reciprocally, DM mice were more submissive compared to single mutants (DM vs 5C: *p* = 0.006, DM vs 2A: *p* = 0.059). Comparison between DM and WT mice did not yield any significant differences. The decreased aggression and increased submission of DM relative to the single mutants were also observed in multiple behavior types, including mounting, chasing for aggression (Supplementary Figure 3A), and cowering and running away for submission (Supplementary Figure 3B). Thus, these results suggest that double mutations alleviate aggressive behavior of both *Kmt2a*-HET and *Kdm5c*-KO mice.

Together, the behavioral studies revealed more pronounced deficits in *Kdm5c*-KO animals compared to *Kmt2a*-HET mice in terms of memory and social interaction, while *Kmt2a*-HET and *Kdm5c*-KO mice shared increased social dominance and aggression. The consequences of double mutations varied between the tests, with clear rescue effects on cognitive tasks, dominant behavior, and aggression, and no effect on social interactions. No behavioral traits were exacerbated in DM. Interestingly, DM mice showed opposite phenotypes in social dominance and aggression compared to single mutants, which is reminiscent of the increased spine density in the DM basolateral amygdala but not in the hippocampal CA1 (Figure 4). These results establish common and unique behavioral deficits of *Kdm5c*-KO and *Kmt2a*-HET mice and mutual suppression between the two genes in some of the traits.

**Figure 4.**
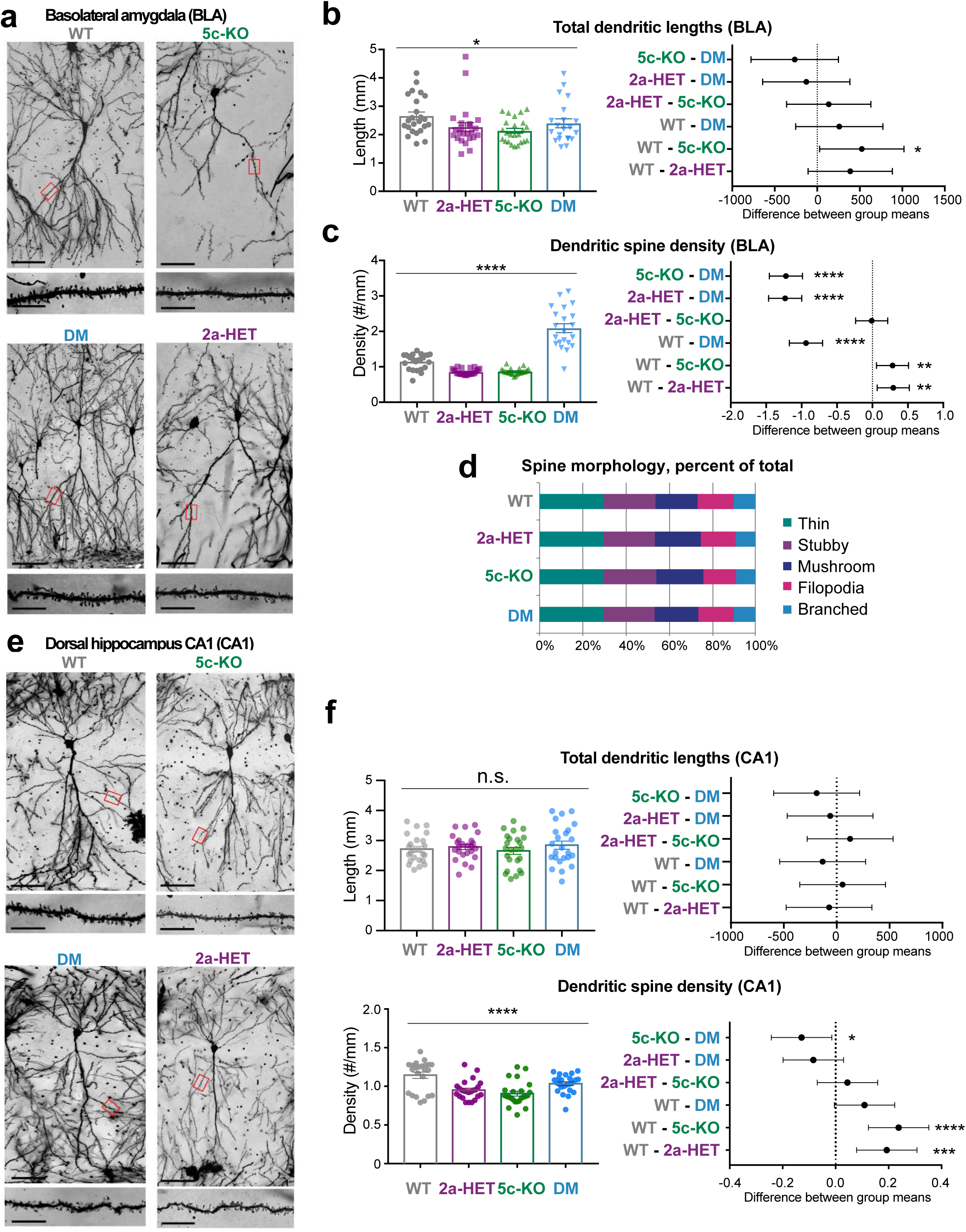
Altered dendrite morphology of *Kdm5c*-KO and *Kmt2a*-HET was reversed in DM animals. (A) Representative images of basolateral amygdala (BLA) pyramidal neurons across all genotypes, depicting overall neuron morphology including dendrite lengths and dendritic spines. Scale bars represent: 100μm (above, whole neuron image), 10μm (below, spine image). **(B and C)** Left panel: Total dendrite lengths **(B)** or spine density **(C)** (mean ± 95% \**p*<0.05, \*\*\*\**p*<0.0001, one-way ANOVA). Right panel: Difference between group means (mean ± 95% confidence intervals, \**p*<0.05, \*\**p*<0.01, \*\*\**p*<0.001, \*\*\*\**p*<0.0001 in Tukey’s multiple comparison test). **(D)** Quantification of spine morphology subtypes represented as percentage of total spines counted. **(E and F)** Morphometry of pyramidal neurons within the dorsal hippocampus CA1. At least 20 neurons from four animals per genotype were quantified for all panels.

### Roles of KMT2A, KDM5C, and their interplay in neuronal morphology

Altered dendrite morphology is a hallmark of many human NDDs, as well as animal models of NDDs (25). We previously found reduced dendritic length and spine density in basolateral amygdala (BLA) neurons of *Kdm5c*-KO adult male mice (21). Assessment of dendritic morphology in *Kmt2a*-HET has not been reported. We first performed comparative dendrite morphometry of pyramidal neurons in the BLA using Golgi staining for the four genotypes (Figure 4). For *Kdm5c*-KO neurons, we recapitulated our previous findings of reduced dendrite lengths (Figure 4A and B, one-way ANOVA followed by Tukey’s multiple comparison test, WT vs 5C: *p* = 0.034) and lower spine density (Figure 4A and C, WT vs 5C: *p* = 0.008). Similar to *Kdm5c*-KO neurons, *Kmt2a*-HET cells exhibited a reduction in spine density (Figure 4B and C, WT vs 2A: p = 0.005) but not in dendritic length (WT vs 2A: p = 0.177).

Dendrite lengths of DM did not differ significantly from WT (Figure 4B, WT vs. DM: *p* = 0.550); however, these lengths were also not different from *Kdm5c*-KO (*Kdm5c*-KO vs. DM: *p* = 0.534), representing a weak restorative effect. In contrast, dendritic spine density of DM showed a significant increase that surpassed a rescue effect (Figure 4C). As morphology of dendritic spines progressively changes during synaptogenesis and development, we also asked whether developmental subtypes of dendritic spines were altered in any genotype. We did not find dramatic changes in spine morphology among the four genotypes (Figure 4D, Supplementary Figure 4), indicating selective requirement of *Kdm5c* and *Kmt2a* for regulation of spine numbers.

Since the ventral hippocampus CA1 (vCA1) receives inputs from BLA (26), we also performed morphometry analyses of pyramidal neurons in this region. Genotype had significant impact on dendritic spine density (one-way ANOVA, F (3, 92) 11.51, p < 1.0 x 10^-4^) but not on dendritic length (F (3, 92) = 0.564, p=0.639). While dendritic length did not show any difference in *Kdm5c*-KO and *Kmt2a*-HET neurons, spine density was decreased in the two single mutants compared to WT (WT vs 2A: *p* = 2.0 x 10^-4^, WT vs 5C: *p* < 1.0 x 10^-4^).

In DM vCA1, spine density showed a trend of degrease compared to WT, yet this did not reach statistical significance (WT vs DM: *p* = 0.066). DM spine density was significantly higher than *Kdm5c*-KO (DM vs 5C: *p* = 0.021) but similar to *Kmt2a*-HET (DM vs 2A: p=0.223). These analyses indicate reduced spine density in both *Kdm5c*-KO and *Kmt2a*-HET vCA1 neurons and its partial correction in DM.

Overall, we conclude that *Kmt2a*-HET and *Kdm5c*-KO share a reduced spine density in both BLA and vCA1. Double mutations led to reversal of spine phenotype in BLA and partial restoration in vCA1, supporting mutually suppressive roles of KMT2A and KDM5C in dendritic spine development.

### Roles of KMT2A and KDM5C in mRNA expression

*Kdm5c*-KO mice exhibit aberrant gene expression patterns in the amygdala and frontal cortex (21), and hippocampus (22). Excitatory-neuron specific conditional *Kmt2a*-KO mice were also characterized with altered transcriptomes in the hippocampus and cortex (19, 20). However, the global gene expression of *Kmt2a*-HET, which is akin to the WDSTS syndrome genotypes, has not been determined. To compare the impact of *Kmt2a*-heterozygosity and *Kdm5c*-KO on the transcriptomes, we performed mRNA-seq using the amygdala and the hippocampus of adult mice with four animals per genotype. We confirmed the lack or reduction of reads from *Kdm5c* exons 11 and 12 and *Kmt2a* exons 8 and 9 (Supplementary Figure 2A-B). The accurate microdissection of brain regions was confirmed by co-clustering of our data with another published mRNA-seq (27) (Supplementary Figure 5A). Principal component analysis (PCA) indicated that transcriptomic divergence was primarily driven by brain regions rather than genotypes (Supplementary Figure 6A)

In the amygdala, we identified 132 differentially expressed genes (DEGs) in *Kdm5c*-KO (5C-DEGs) and no 2A-DEG (*padj* < 0.1, n =4, Figure 5A and Supplementary Figure 6). The hippocampus yielded a consistently higher number of DEGs across the mutant genotypes (344 5C-DEGs and 4 2A-DEGs). The small number of 2A-DEGs is likely due to the one remaining copy of *Kmt2a*. Given increased social dominance (Figure 3B) and clear reduction of dendritic spines (Figure 4) in *Kmt2a*-HET mice, we reasoned that such phenotypes might be associated with mild gene misregulation, which was not detected by DEseq2. To be able to analyze the gene expression in *Kmt2a*-HET, we set a relaxed cut off (*p* < 0.005) and retrieved 78 and 139 genes as 2A-DEG from the amygdala and hippocampus, respectively (Figure 5B and Supplementary Figure 6). We found an overall similarity in gene misregulation between the two brain regions for both *Kdm5c*-KO and *Kmt2a*-HET (Supplementary Figure 7A-B). Table S1 lists all DEGs found in the study.

**Figure 5.**
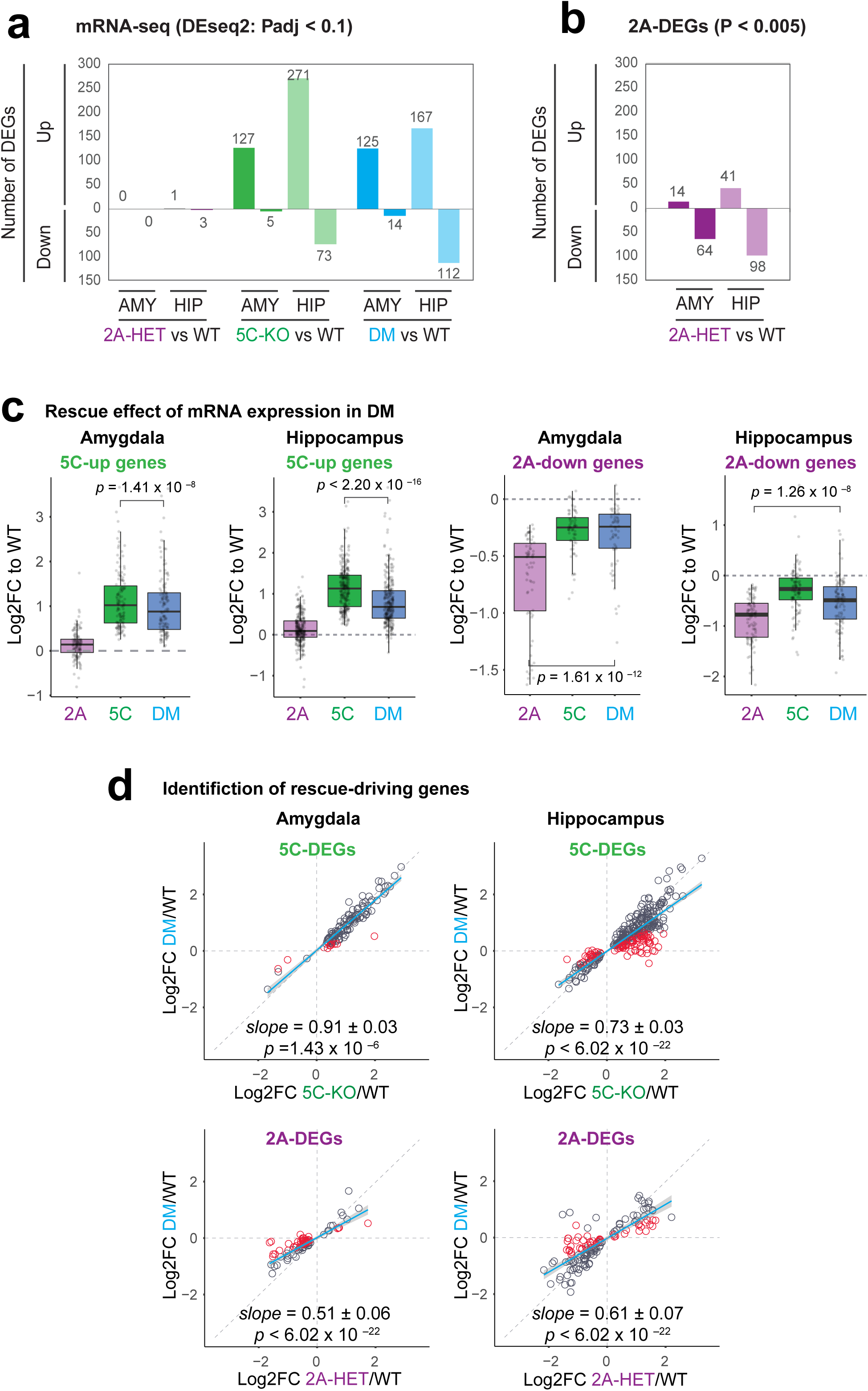
The transcriptomes in the amygdala and hippocampus. (A) Number of differentially expressed (DE) genes across genotypes were determined using a threshold of *padj*-value < 0.1. or relaxed cut-off of *p* < 0.005 for *Kmt2a*-HET in **(B)** (see also Table S1). We analyzed amygdala and hippocampal tissues from four animals for each genotype. **(C)** Behavior of single-mutant DEGs in DM. Log2 fold change of DEGs relative to WT were plotted across the three mutants. Boxplot features: box, interquartile range (IQR); bold line, median; gray dots, individual genes. Associated *p* values result from Wilcoxon signed-rank tests. **(D)** Identification of rescue-driving genes and regression analysis. Blue fitting lines and slopes result from linear regression of log2 fold changes between the two genotypes. Gray shade: 95% confidence interval. *P*-values indicate probability of the null hypothesis that the fitting line does not differ from 1 (48). Red circles: rescue-driving genes (see method).

We next compared the gene expression profiles between the single mutants. The majority of 5C-DEGs were upregulated and 2A-DEGs were downregulated (Figure 5A and 5B, Supplementary Figure 6). This result agrees with KMT2A’s primary role as a transcriptional coactivator and KDM5C’s suppressive activity on transcription by removing this mark, yet KDM5C can also act as a positive regulator of transcription (28). If KMT2A and KDM5C counteract, DEGs would be oppositely misregulated between the single mutants. Indeed, we found that 28-29% of 5C-DEGs and 8-17% of 2A-DEGs show signs of opposite regulation in the other single mutant (Supplementary Figure 8A-B). Substantial fractions of DEGs were misregulated in the same directions (Supplementary Figure 8A-B), which might be due to indirect consequences of gene manipulations and altered circuits.

### Impact of double mutations on mRNA expression

If the double mutations fully restore the normal transcriptome, the DM transcriptome should resemble to that of WT over single mutants. However, the DM brain tissues had a similar number of DEGs as *Kdm5c*-KO (Figure 5A). Many DM-DEGs overlap with 5C- DEGs (84 in the amygdala and in the hippocampus), and 41 DM-DEGs overlap with 2A- DEGs in the hippocampus (Supplementary Figure 8C). Thus, the DM transcriptome still retains some mRNA misregulations of single mutants and does not fully return to the normal state.

To assess the rescue effect more quantitatively, we compared fold changes of single-mutant DEGs as a group between genotypes. Expression of 5C-up-regulated genes was significantly lower in the DM amygdala and hippocampus (Paired Wilcoxon signed-rank test, *p* < 1.41 x 10^-8^, Figure 5C). Reciprocally, expression of 2A-down-regulated genes was significantly higher in DM brain tissues (*p* < 1.26 x 10^-8^, Figure 5C). We further examined how individual DEGs behaved in DM. The fold changes of 5C-DEGs and 2A-DEGs in single mutants clearly correlated with those in DM, indicating that single-mutant DEGs are similarly dysregulated in DM in general. However, the slopes of linear regression were lower than 1 with statistical significance, suggesting the partial rescue effect (Figure 5D). The mildest rescue effect was observed in 5C-DEGs in the DM amygdala (linear regression slope=0.91±0.03), while 5C-DEGs in the hippocampus and 2A-DEGs-R exhibit more pronounced rescue effects (slope: 0.51∼0.73). These results indicate that misregulation of genes in both *Kdm5c*-KO or *Kmt2a*-HET was partially corrected in the DM brain tissues.

We then sought to isolate genes that might have contributed to the rescue effects at cellular and behavioral levels. As shown by the red circles in Figure 5D, some DEGs exhibit stronger rescue effects than others. We selected these genes as potential drivers of rescue effects (see Materials & Methods). While 10 genes in the amygdala and 81 genes in the hippocampus drove rescue effects of 5C-DEGs, 36 genes in the amygdala and 46 genes in the hippocampus contributed to the partial restoration of normal 2A-DEGs expression in DM (Supplementary Data 2). We then performed pathway analysis on these rescue-driving genes using the Enrichr program. While 2A- rescue genes did not yield any statistically significant enrichment (*padj* < 0.05), 5C- rescued genes showed an enrichment of a mouse KEGG pathway, “neuroactive ligand-receptor interaction” (*padj* = 5.60 x 10^-3^, Odds ratio: 6.01). The following 10 genes contributed to the enrichment: *Pomc*, *Adcy2*, *Agt*, *Adora2a*, *Mc3r*, *Gabre*, *P2ry14*, *Tacr1*, *Trh*, *and Lhb*. Except for *P2ry14*, all these genes have known roles in learning and memory or aggression (Table 1). Many of these genes were expressed at low levels in the WT and were derepressed following the KDM5C loss (Supplementary Data 2), suggesting roles for KDM5C in suppressing aberrant gene expression.

**Table 1.**
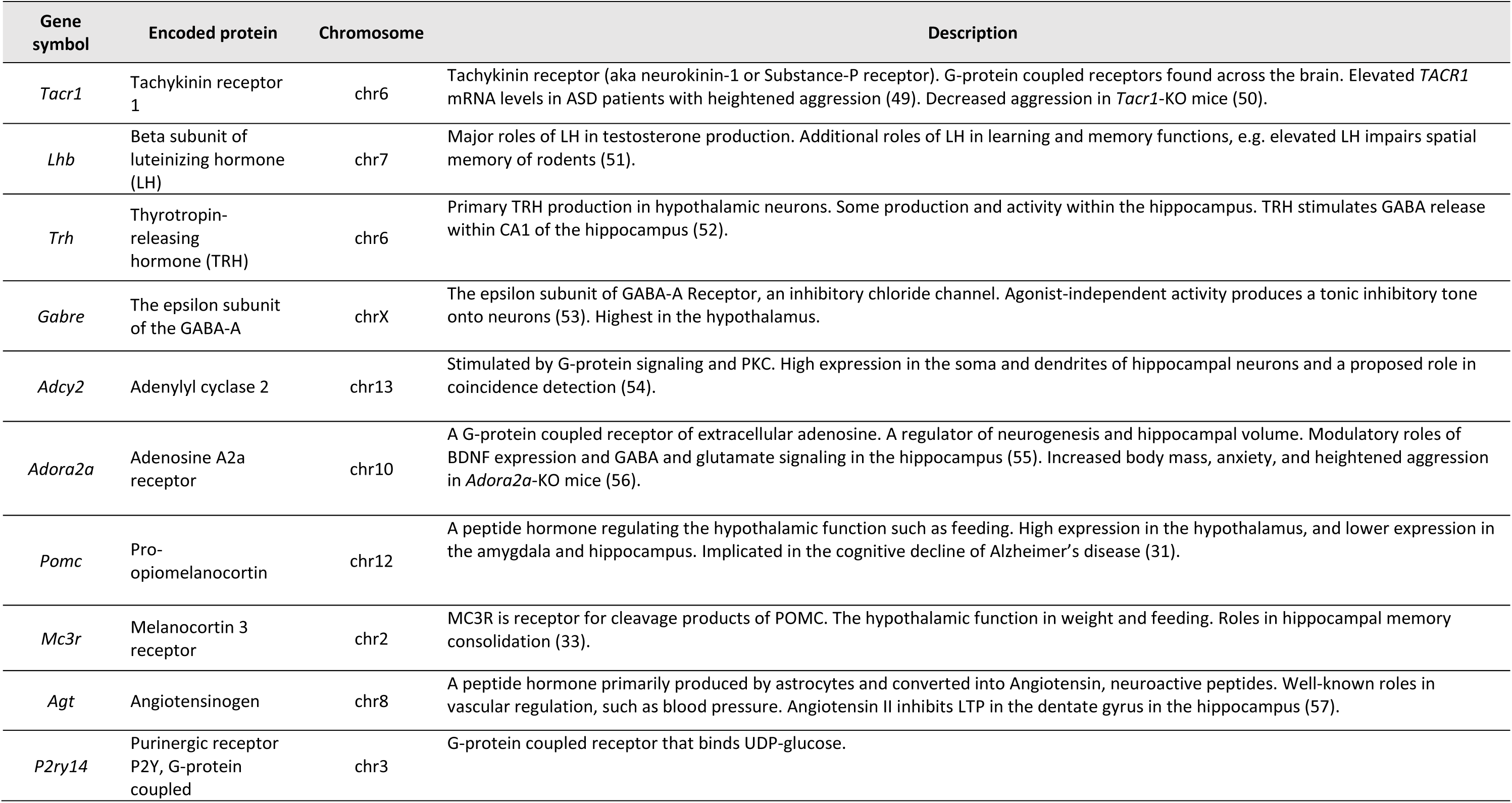
Rescued 5C-DEGs that drove enrichment of the KEGG pathway “Neuroactive ligand-receptor interaction”

Interestingly, most of these genes encode peptide hormones, their receptors, or downstream signaling molecules. For example, Pro-opiomelanocortin (POMC) is a peptide hormone primarily known for its roles in regulating hypothalamic functions, such as feeding (29). In addition to its high expression in the hypothalamus, *Pomc* is also expressed in the amygdala and the hippocampus (30). POMC signaling in these regions has been implicated in the cognitive decline of Alzheimer’s disease (31). Another gene in the list, *Mc3r*, encodes a POMC receptor. *Mc3r* is expressed in the hypothalamus and limbic structures such as the amygdala and hippocampus (32). Similar to *Pomc*, *Mc3r*’s roles have been primarily studied in the hypothalamus, yet this gene has also been implicated in hippocampal memory consolidation (33). Collectively, these results raise the possibility that aberrant peptide hormone signaling and its normalization underlie phenotypic outcomes of *Kdm5c*-KO and DM mice.

### H3K4me3 landscapes in WDSTS and MRXSCJ models

We sought to determine the impact of KMT2A- and KDM5C-deficiencies and double mutation on the H3K4me3 landscape within the amygdala. In Western blot analyses, global H3K4me1-3 levels were not altered dramatically in any mutant (Figure 6A, Supplementary Figure 9). We thus performed H3K4me3 chromatin immunoprecipitation coupled with deep sequencing (ChIP-seq) to probe local changes genome-wide. To assess the IP specificity, we spiked-in an array of recombinant nucleosomes carrying 15 common methylations along with DNA barcodes appended to the Widom601 nucleosome positioning sequence (34) (see Methods). The two recombinant H3K4me3 nucleosomes dominated the Widom601-containing DNA in all IP reactions with negligible signals from other methylation states such as H3K4me1 or H3K4me2 (Supplementary Figure 10A), demonstrating a superb specificity of the ChIP.

**Figure 6.**
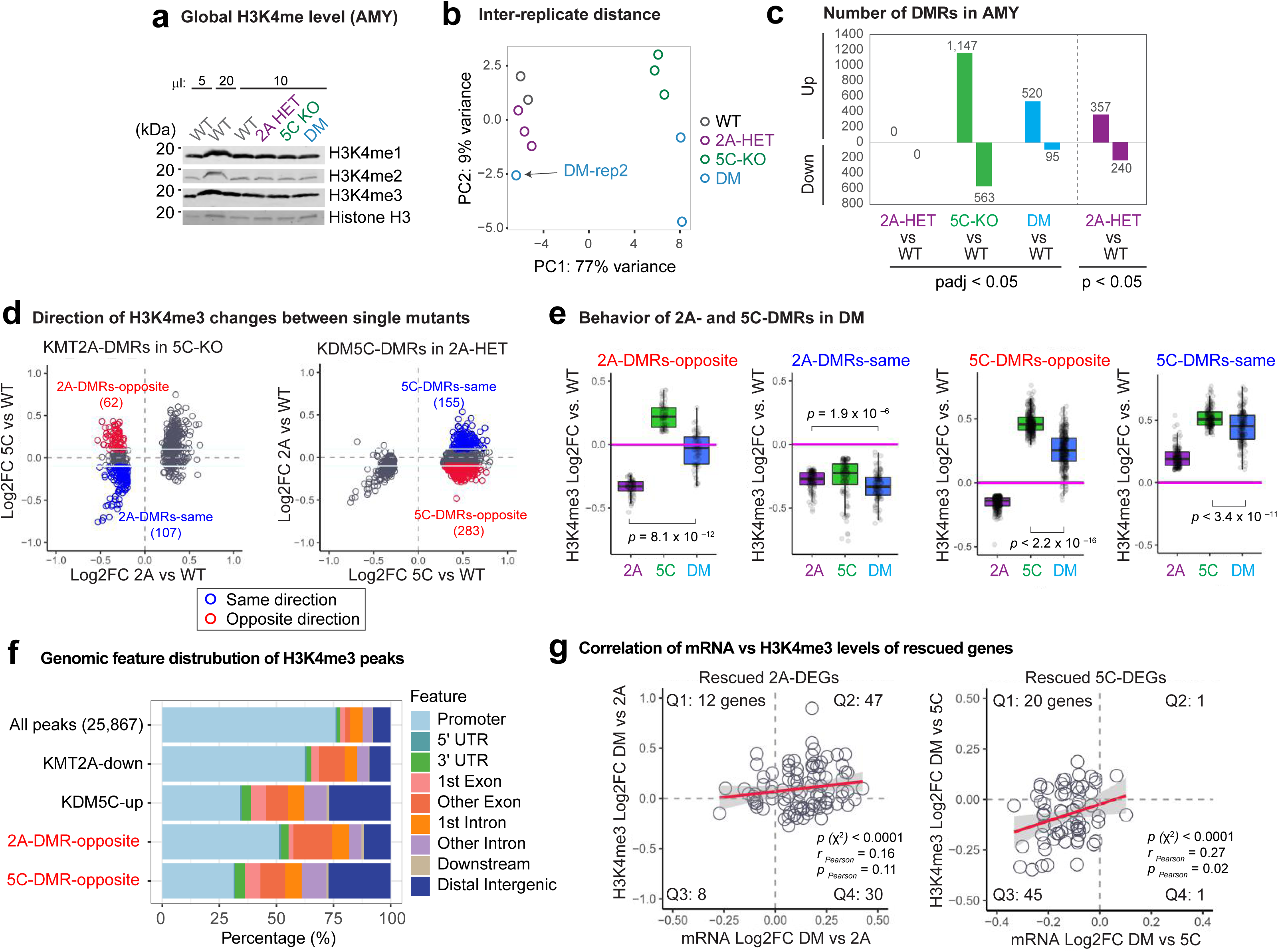
Altered H3K4me3 landscapes in the amygdala and rescue effect in DM. (A) Western blot of whole brain lysates showing unchanged global H3K4 methylation across genotypes. Total histone H3 was detected using an antibody recognizing the C-terminus of H3, and used as a control for equal loading. **(B)** PCA analysis of H3K4me3 ChIPseq replicates. We analyzed amygdala tissue from 2 to 3 animals (rep1-3) for each genotype (see Methods). **(C)** Number of H3K4me3 DMRs in 2a-HET, 5c-KO, or DM compared to WT across the genome. 2A-HET DMRs are retrieved with a relaxed threshold (*p* < 0.05). **(D)** Direction of H3K4me3 changes between single mutants. Genes are colored based on the direction of misregulation between the single mutants. **(E)** Behavior of single-mutant DMRs in DM. Log2 fold change of DMRs relative to WT were plotted across the three mutants. Boxplot features: box, interquartile range (IQR); bold line, median; gray dots, individual genes. Associated *p* values result from Wilcoxon signed-rank tests. **(F)** Genomic features of H3K4me3 peaks. **(G)** Involvement of H3K4me3 restoration near rescued DEGs. H3K4me3 log2 fold changes (FC) are plotted as function of mRNA log2 FC between DM and single mutants. Red line: linear regression fitting line with 95% confidence interval (gray shade). The gene numbers fall into each of the four quadrants as indicated (Q1-Q4). Results (*p*) of χ^2^ test and Pearson’s correlation coefficient (*r*) and p-value (*p*) are noted.

PCA analyses of H3K4me3 distribution indicated that WT and *Kmt2a*-HET are close each other, while *Kdm5c*-KO and DM cluster separately from the WT-2a-HET cluster. Reminiscent of the transcriptome data (Figure 5), we did not recover any differentially methylated regions (DMRs) in *Kmt2a*-HET, while *Kdm5c*-KO had 1,147 hypermethylated regions (Figure 6C, Supplementary Figure 9, padj < 0.05, see method). In *Kdm5c*-KO, 583 loci are hypomethylated compared to WT, and these regions are highly methylated regions in WT (Figure 6C and Supplementary Figure 10). By relaxing the threshold, we obtained weak 2A-HET DMRs (Figure 6C, 357 hypermethylated and 240 hypomethylated). We then examined how these single-mutant DMRs behaved in the other single mutant. Compared to the transcriptome data (Supplementary Figure 8), we were able to recover more DMRs that are misregulated in opposite directions between the single mutants (Figure 6D). These results suggest that, as a primary substrate of the two enzymes, H3K4me3 changes represent the action of KMT2A and KDM5C more directly compared to mRNA expression. Comparable numbers of DMRs still show same-direction H3K4me3 changes, which may again reflect indirect consequences of gene manipulations.

### Rescue effect of H3K4me3 landscapes in DM

We next examined the impact of double mutations on H3K4me3 distributions. Notably, in the PCA plot, the second replicate of DM showed a stronger rescued compare to other replicates, indicating that expressivity of rescue effect varies among individual animals (Figure 6B). The number of DMRs in DM are smaller than those of single mutants, indicating a global rescue of aberrant H3K4me3 (Figure 6C). We then assessed how single-mutant DMRs behaved in DM. Note that, we focus on hypomethylated loci in *Kmt2a*-HET and hypermethylated loci in *Kdm5c*-KO (colored loci in Figure 6D) based on their enzymatic activities. The opposite-direction 2A-DMRs showed complete normalization in DM, while the same-direction 2A-DMRs were even more hypomethylated in DM (Figure 6E). Likewise, the opposite-direction 5C-DMRs showed more pronounced rescue effect in DM compared to same-direction 5C-DMRs (Figure 6E). This direction-dependent rescue effect persisted even when we omitted DM rep2 from the analysis (Supplementary Figure 11). These results strongly support the idea that KMT2A and KDM5C counteract to normalize H3K4me3 levels in specific genomic loci.

Peak annotation revealed that rescued DMRs tend to be outside the promoters, such as distal intergenic regions, introns, and internal exons (Figure 6F). ChIP-seq tracks of exemplary loci are represented in Supplementary Figure 12. Finally, we tested whether the rescue effects in gene expression involve normalization of H3K4me3. We took all rescued 2A-DEGs and plotted DM vs 2A fold changes of H3K4me3 as a function of DM vs 2A mRNA changes (Figure 6G). Rescued 2A-DEGs mostly showed an increase in mRNA levels in DM (Q2 and Q4). When mRNA expression increases from 2A-HET to DM, H3K4me3 levels also increase, which results in the larger number of genes within the upper right quadrant of the plot (Q2, 47 genes, *p* < 1.0 x 10^-4^, χ^2^ test). Similarly, when mRNA expression decreases from 5C-KO to DM, the majority of genes are accompanied by decreased H3K4me3 nearby (Q3: 45 genes vs Q1: 20 genes, *p* < 1.0 x 10^-4^, χ^2^ test). Furthermore, the magnitude of mRNA and H3K4me3 changes show positive correlations (Pearson *r* = 0.16, and *r* = 0.27 for 2A- and 5C-DEGs, respectively). Thus, misregulation of mRNA expression and its partial correction involve corresponding restoration of the H3K4me3 levels.

## Discussion

The present work, to our knowledge, represents the first genetic interactions between mammalian methyl histone writer and eraser *in vivo*. Interplay of opposing chromatin-modifying enzymes has been characterized in several biological processes and species. For example, DNMT3A DNA methyltransferase and TET2 dioxigenase, which oppose over DNA CpG methylation, can both counteract and cooperate to regulate gene expression in hematopoietic stem cells (35). Set1 and Jhd2, the sole H3K4me writer-eraser pair in yeast, have been shown to co-regulate transcription (36). However, a fundamental question remained — is there any specific writer-eraser pairing in highly duplicated gene families for a single chromatin modification? Mishra et al. showed that KDM5A antagonizes KMT2A, KMT2B, and KMT2D to modulate the transient site-specific DNA copy number gains in immortalized human cells (37). Cao et al. found that failed differentiation of mouse embryonic stem cells due to *Kmt2d* deletion can be rescued by *Kdm1a* knockdown (38). These pioneering efforts identified functional interplay between the opposing enzymes *in vitro*; however, no *in vivo* study has been reported. Thus, the present study substantially advances our understanding of how methyl-histone enzymes functionally interact.

Brain development is particularly relevant to the H3K4me dynamics, because a cohort of neurodevelopmental disorders has been genetically associated with impaired functions of these enzymes, as discussed earlier. Unlike previous studies using chemical approaches that block multiple chromatin regulators (13–15), we demonstrated that manipulation of a single enzyme, KMT2A or KDM5C, is sufficient to reverse many neurological traits in both of the two single-mutant models. This study motivates interrogations of human populations to test if damaging mutations in writer and eraser enzymes can coincide in unaffected individuals. Our work also opens a new avenue for future studies to delineate the full interplay between the H3K4me-regulatory enzymes.

Several challenges remain, especially in linking molecular functions of KMT2A and KDM5C to cellular and behavioral outcomes. First, we measured gene expression and H3K4me3 in adult brain tissues, a mixture of many neuron and glia types, which may mask potentially important molecular changes. Increasing spatiotemporal resolution of the molecular study is an important future direction. Second, histone modifying enzymes, including KDM5C, can exert non-enzymatic function (11, 39). Although it has not been reported, there might be a non-histone substrate for KMT2A and KDM5C, as shown in other histone modifying enzymes (12). Thus, our study does not rule out the impact of these non-canonical roles of the two enzymes. Finally, in a fly model of KMD5-intellctual disability (40, 41), impaired intestinal barrier and the altered microbiota contribute to abnormal social behavior, pointing to a non-tissue-autonomous mechanism (42). Future studies are warranted to isolate the causal events for observed rescue effects. The first step would be to test the functional contribution of candidate genes, many of which encode peptide-hormone signaling factors (Table 1).

Increased social dominance is a novel behavioral trait we observed in both WDSTS and MRXSCJ mouse models. The amygdala is well known to mediate social behaviors. For example, lesions of BLA result in decreased aggression-like behavior and increased social interactions, and changes in transcriptional regulation in BLA are observed after social interactions (43). The dorsal hippocampus CA1 is a direct recipient region of BLA inputs (26). Decreased BLA and CA1 spine densities in *Kmt2a*-HET and *Kdm5c*-KO mice inversely correlate with increased social dominance and aggression (Figures 3 and 4). Together, these observations imply that decreased spine density is not a cause of increased social dominance, but rather reflects compensatory reduction of net synaptic strength due to increased excitation and/or a loss of inhibitory control in this circuit. Thus, determining the connectivity of the amygdala with other regions, including prelimbic, infralimbic, and orbitofrontal cortices as well as the ventral hippocampus, will be critical for understanding the changes in social behaviors in both WDSTS and MRXSCJ models.

It is important to note that the double mutations introduced in our mice were constitutive, and therefore a lifetime of adaptation to loss of these two major chromatin regulators may occur from early developmental stages. A more realistic therapeutic strategy may be acute inhibition of KDM5C and KMT2A in the juvenile or mature brain. Previous work characterizing mouse models with excitatory-neuron specific ablation of *Kdm5c* or *Kmt2a* via *CamKII*-Cre found that conditional *Kmt2a* deletion led to clear learning deficits (20), while cognitive impairments in the conditional *Kdm5c-*KO mice were much milder than those of constitutive *Kdm5c-*KO mice (22). These results suggest a developmental origin of phenotypes in *Kdm5c*-KO. Future investigations are needed to address whether the effects of acute inhibition of opposing enzymes in these mouse models can restore such neurodevelopmental deficits.

## Materials & methods

### Statistics and Reproducibility

The proposed study was conducted with varied numbers of individual animals depending on the experiments. Data acquisition and analysis were conducted blindly except for molecular measurements including Western blots and genomics analyses. We did not remove any particular data points for their acquisition or analyses. Statistical tests were chosen based on the distribution of the data. Details of statistics and sampling are outlined below in each section.

### Mouse

The *Kdm5c*-KO allele was previously described and maintained in C57BL/6J congenic background (21). *Kmt2a*-HET mice were generated by crossing previously described *Kmt2a*-flox (exons 8 and 9) mice with B6.129-Gt(ROSA)26Sor^tm1(cre/ERT2)Tyj^/J-Cre mice (44). Our strategy was to use F1 hybrid for all studies as previously recommended as a standard practice to eliminate deleterious homozygous mutations, which can result in abnormalities in the congenic lines, e.g. deafness in C57/BL6 (45). We backcrossed *Kmt2a*^+/-^ mice onto the desired 129S1/SvImJ strain, by marker-assisted accelerated backcrossing through Charles River Labs. *Kmt2a*^+/-^ mice were bred to the N4 generation at minimum, where mice were >99% congenic for 129S1/SvImJ. All experimental mice were generated as F1 generation hybrids from mating between 129S1/SvImJ *Kmt2a*^+/-^ males and C57Bl/6 *Kdm5c*^+/-^ females: WT males (*Kmt2a*^+/+^, *Kdm5c*^+/y^); *Kdm5c*-KO males (*Kmt2a*^+/+^, *Kdm5c*^-/y^); *Kmt2a*-HET males (*Kmt2a^+/-^*, *Kdm5c*^+/y^); and *Kdm5c-Kmt2a*- DM males (*Kmt2a*^+/-^, *Kdm5c*^-/y^). Genotypes were confirmed using the following primers: for *Kmt2a*, 5’-GCCAGTCAGTCCGAAAGTAC, 5’-AGGATGTTCAAAGTGCCTGC, 5’- GCTCTAGAACTAGTGGATCCC; for *Kdm5c*, 5’-CAGGTGGCTTACTGTGACATTGATG, 5’-TGGGTTTGAGGGATACTTTAGG, 5’-GGTTCTCAACACTCACATAGTG.

All mouse studies complied with the protocols (PRO00008568: Iwase and PRO00008807: Tronson) by the Institutional Animal Care & Use Committee (IACUC) of The University of Michigan.

### Western blot analysis

Total proteins from adult brain tissues were subjected to Western blot analysis using in-house anti-KDM5C (21) and anti-GAPDH antibodies (G-9, Santa Cruz). For histone proteins, nuclei were enriched from the Dounce-homogenized brain tissues using Nuclei EZ prep Kit (Sigma, NUC-101). DNA were digested with micrococcal nuclease (MNase, NEB) for 10 minutes at room temperature, and total nuclear proteins were extracted by boiling the samples with the SDS-PAGE sample buffer. The following antibodies were used for Western blot analyses: anti-H3K4me3 (Abcam, ab8580, 1:1000), anti-H3K4me2 (Thermo, #710796, 1:1000), anti-H3K4me1 (Abcam, ab8895, 1:1000), and anti-H3 C-terminus (Millipore, CS204377, 1:2000).

### Brain histology

Mice were subjected to transcardial perfusion according to standard procedures. Fixed brains were sliced on a freeze microtome, yielding 30 μm sections that were then fixed, permeabilized, blocked, and stained with DAPI. Slides were imaged on an Olympus SZX16 microscope, with an Olympus U-HGLGPS fluorescence source and Q Imaging Retiga 6000 camera. Images were captured using Q-Capture Pro 7 software. Data were collected in a blind fashion, where samples were coded and genotypes only revealed after data collection was complete.

### Behavioral paradigms

Prior to behavioral testing, mice were acclimated to the animal colony room for one week single-housing in standard cages provided with a lab diet and water *ad libitum*. A 12-hour light-dark cycle (7:00AM-7:00PM) was utilized with temperature and humidity maintained at 20 ±2 °C and >30%, respectively. The University of Michigan Committee on the Use and Care of Animals approved all tests performed in this research. Five tests, listed in order of testing, were performed: Novel Object Recognition (five days), Context Fear Conditioning (two days), Three-Chambered Social Interaction (two days), Social Dominance Tube Test (three to four days), and Resident-Intruder (two to three days). All testing was conducted in the morning by experimenters blind to genotype. The cleaning agent used in every test between each trial was 70% ethanol. Data were collected in a blind fashion, where mice were coded and genotypes were only revealed after testing was complete.

#### Contextual Fear Conditioning

Mice were placed into a distinct context with white walls (9 ¾ × 12 ¾ × 9 ¾ in) and a 36 steel rod grid floor (1/8 in diameter; ¼ spaced apart) (Med-Associates, St. Albans, VT) and allowed to explore for three minutes, followed by a two-second 0.8 mA shock, after which mice were immediately returned to their home cages in the colony room. Twenty-four hours later, mice were returned to the context and freezing behavior was assessed with NIR camera (VID-CAM-MONO-2A) and VideoFreeze (MedAssociates, St Albans, VT). Freezing levels were compared between genotypes using a between-groups analysis (one-way ANCOVA) with genotype as the between-subjects factor, and “cohort” as the covariate, to reduce variability as a result of multiple cohorts. Planned comparisons between genotypes were conducted, with Bonferroni correction for multiple comparisons.

#### Novel Object Recognition

Mice were first habituated to testing arenas (40 x 30 x 32.5 cm^3^) in three, 10-minute sessions over six consecutive days. Twenty-four hours later, mice were allowed to explore two identical objects (a jar or egg, counterbalanced across animals) for two, 10-minute trials spaced three hours apart. All animals were returned to the arena, tested 24 hours after the first training session, and presented with one training object (“familiar” object: jar or egg) and one “novel” object (egg or jar). Exploration of the objects was defined as nose-point (sniffing) within 2 cm of the object. Behavior was automatically measured by Ethovision XT9 software using a Euresys Picolo U4H.264No/0 camera (Noldus, Cincinnati, OH). Preference was calculated as the time spent exploring novel object/total time exploring both objects. One-sample Wilcoxon signed-rank tests against 50% (no preference) was used to establish whether animals remembered the original objects.

#### Three-Chambered Social Interaction

Mice were placed into a three-chambered apparatus consisting of one central chamber (24 x 20 x 30 cm^3^) and two identical side chambers (24.5 x 20 x 30 cm^3^) each with a containment enclosure (8 cm diameter; 18 cm height; grey stainless steel grid 3 mm diameter spaced 7.4 mm apart) and allowed to habituate for 10 minutes. Twenty-four hours later, mice were returned to the apparatus that now included a 2-3 month-old stranger male mouse (C57BL/6N) on one side of the box (“stranger”), and a toy mouse approximately the same size and color as the stranger mouse on the other (“toy”). Exploration of either the stranger or toy was defined as nose-point (sniffing) within 2 cm of the enclosure and used as a measure of social interaction. Behavior was automatically scored by Ethovision XT9 software as described above, and social preference was defined as time exploring stranger/total time spent exploring (stranger + toy). One-sample Wilcoxon signed-rank tests against 50% (no preference) was used to establish whether animals remembered the original objects. Differences between genotypes were analyzed using an ANCOVA with cohort as a covariate. Planned comparisons with Bonferroni correction for multiple comparisons were used to assess differences between genotypes.

#### Social Dominance Tube Test

Twenty-four hours prior to testing, mice were habituated to a plastic clear cylindrical tube (1.5 in diameter; 50 cm length) for 10 minutes. During the test, two mice of different genotypes were placed at opposite ends of the tube and allowed to walk to the middle. The match concluded when the one mouse (the dominant mouse) forced the other mouse (the submissive mouse) to retreat with all four paws outside of the tube (a “win” for the dominant mouse). Each mouse underwent a total of three matches against three different opponents for counterbalancing. Videos were recorded by Ethovision XT9 software as described above, and videos were manually scored by trained experimenters blind to genotype. The number of “wins” was reported as a percentage of total number of matches. Data were analyzed using an Exact Binomial Test with 0.5 as the probability of success (win or loss).

#### Resident-Intruder Aggression

Resident-intruder tests were used to assess aggression. Tests were performed on consecutive days, where the resident mouse was exposed to an unfamiliar intruder mouse for 15 minutes. A trial was terminated prematurely if blood was drawn, if an attack lasted continuously for 30 seconds, or if an intruder showed visible signs of injury after an attack. Resident mice were assessed for active aggression (darting, mounting, chasing/following, tail rattling, and boxing/parrying), as well as submissive behaviors (cowering, upright, running away). Intruder mice were assessed for passive defense (freezing, cowering, and digging). Behavior was recorded and videos scored manually by experimenters blind to genotype. A repeated measures analysis, with cohort as a covariate, was used for each aggressive (genotype x aggression measures ANOVA) and submissive behavior (genotype x submissive) to analyze aggressive behaviors. Planned comparisons for genotype, with Bonferroni corrections for multiple comparisons were used to further analyze specific effects of genotype.

### Neuronal Golgi staining and morphological analyses

Brains from adult (2-8 months) mice were dissected and incubated in a modified Golgi-Cox solution for two weeks at room temperature. The remaining procedure of Golgi immersion, cryosectioning, staining and coverslipping was performed as described previously (21). Four animals were used for each genotype, and pyramidal neurons in the basolateral amygdala and dorsal hippocampus CA1 per animal were quantified: N=24 neurons for WT, *Kmt2a*-HET and *Kdm5c*-KO, and N=27 neurons for DM. Quantification was done using commercially available software, NeuroLucida (v10, Microbrightfield, VT), installed on a Dell PC workstation that controlled a Zeiss Axioplan microscope with a CCD camera (1600 x 1200 pixels) and with a motorized X, Y, and Z focus for high-resolution image acquisition (100X oil immersion) and quantifications. The morphological analyses included: dendritic lengths, spine counts, and spine subtype morphology. All sample genotypes were blinded to the analysts throughout the course of the analysis.

The criteria for selecting candidate neurons for analysis were based on: (1) visualization of a completely filled soma with no overlap of neighboring soma and completely filled dendrites, (2) the tapering of most distal dendrites; and (3) the visualization of the complete 3-D profile of dendritic trees using the 3-D display of the imaging software.

For quantitative analysis of spine subtypes (thin, stubby, mushroom, filopodia, and branched spines), only spines orthogonal to the dendritic shaft were included in this analysis, whereas spines protruding above or beneath the dendritic shaft were not sampled. This principle remained consistent throughout the course of analysis.

After completion, the digital profile of neuron morphology was extrapolated and transported to a multi-panel computer workstation, then quantitated using NeuroExplorer program (Microbrightfield, VT), followed by statistical analysis (one- and two-way ANOVAs, *p* < 0.05).

### RNA-seq

Brains from adult (4.5 to 8 months) male mice were microdissected to obtain the amygdala and hippocampus from Bregma ∼ 4.80 mm regions. N=4 animals were used per genotype. The ages of mice used for genomics study, RNA-seq and ChIP-seq, are summarized in Supplementary Data 3. Tissue was homogenized in Tri Reagent (Sigma). Samples were subjected to total RNA isolation, and RNA was purified using RNEasy Mini Kit (Qiagen). ERCC spike-in RNA was added at this stage, according to manufacturer’s instructions (Life Technologies). Libraries were prepared using the NEBNext® Ultra™ II Directional RNA Library Prep Kit with oligo-dT priming. Multiplexed libraries were pooled in approximately equimolar ratios and purified using Agencourt RNAClean XP beads (Beckman Coulter).

Libraries were sequenced on the Illumina Novaseq 6000 platform, with paired-end 150 base pair reads (24-35 million reads/library), according to standard procedures. Reads were mapped to the mm10 mouse genome (Gencode) using STAR (v2.5.3a), where only uniquely mapped reads were used for downstream analyses. Duplicates were removed by STAR, and a counts file was generated using FeatureCounts (Subread v1.5.0). BAM files were converted to bigwigs using deeptools (v3.1.3). Differentially expressed (DE) genes were called using DESeq2 (v1.14.1). Data analyses were performed with RStudio (v1.0.136). Fold change heatmaps was created using gplots heatmap.2 function.

To validate the microdissection of hippocampus and amygdala, we compared our RNA-seq datasets with similar RNA-seq data from Arbeitman (27) datasets, which involved the cerebellum, cortex, hippocampus, and amygdala. Briefly, count data underwent variance stabilizing transformation via DEseq2 vst function and Euclidean distances of the transformed values were calculated by dist command and the heatmap was generated by the pheatmap function. Linear regression analysis was performed using the smtr v3.4-8 slope.test function with intercept = FALSE and robust=FALSE options. Rescue-driving genes were chosen as genes that satisfy two conditions using R, **1)** abs (log2FC (single mutant vs DM) > 2, **2)** abs (log2FC (DM vs WT) < 0.7.

### ChIP-seq

Amygdala tissues were microdissected from adult (8-14.5 months) male mice. N=2 animals were used for WT, and N=3 animals were used for *Kmt2a*-HET, *Kdm5c*-KO, and DM as biological replicates. Nuclei were isolated using a Nuclei EZ prep Kit (Sigma, NUC-101), and counted after Trypan blue staining. Approximately 20,000 nuclei for each replicate were subjected to MNase digestion as previously described (46). We essentially followed the native ChIP-seq protocol (46) with two modifications. One was to use a kit to generate sequencing libraries in one-tube reactions (NEB, E7103S). Another modification was to spike-in the panel of synthetic nucleosomes carrying major histone methylations (EpiCypher, SKU: 19-1001) (34). For ChIP, we used the rabbit monoclonal H3K4me3 antibody (Thermo, clone #RM340, 2 μg).

Libraries were sequenced on the Illumina NextSeq 500 platform, with single-end 75 base-pair sequencing, according to standard procedures. We obtained 20-59 million reads per sample. Reads were aligned to the mm10 mouse genome (Gencode) and a custom genome containing the sequences from our standardized, synthetic nucleosomes (EpiCypher) for normalization, using Bowtie allowing up to two mismatches. Only uniquely mapped reads were used for analysis. The range of uniquely mapped reads for input samples was 38-44 million reads. All IP replicates had a mean of 9.1 million uniquely mapped reads (range: 7.4-13.9 million). The enrichment of mapped synthetic spike-in nucleosomes compared to input was calculated and used as a normalization coefficient for read depth of each ChIP-seq replicate.

Peaks were called using MACS2 software (v 2.1.0.20140616) using input BAM files for normalization, with filters for a q-value < 0.1 and a fold enrichment greater than 1. Common peak sets were obtained via DiffBind, and count tables for the common peaks were generated with the Bedtools multicov command. We removed “black-list” regions that often give aberrant signals. Resulting count tables were piped into DEseq2 to identify DMRs incorporating the synthetic nucleosome normalization into the read depth factor. We used ChIPseeker to annotate H3K4me3 peaks with genomic features including mm10 promoters (defined here as ±1 kb from annotated transcription start site [TSS]). Normalized bam files were converted to bigwigs for visualization in the UCSC genome browser. Genes near peaks were identified by ChIPseeker. RNA-seq and ChIP-seq data were integrated using standard R commands and rescued amygdala DEGs were chosen as genes that meet two criteria, **1)** abs (log2FC (single mutant vs DM) > 1, **2)** abs (log2FC (DM vs WT) < 0.7. All scripts used in this study are available upon request.

### Data Availability

The RNA-seq and ChIP-seq are available in NCBI’s Gene Expression Omnibus (47). Accession numbers are GSE127722 for RNA-seq, GSE127817 for ChIP-seq and GSE127818 for SuperSeries.

## Acknowledgements

We thank Dr. Ken Kwan, Mandy Lam, and Own Funk for their assistance with the RNA-seq library preparation protocol and use of their microscope; Chris Gates for his assistance with RNA-seq analyses; and Clara Farrehi, Jordan Rich, and Demetri Tsirukis for their assistance with experiments for transcriptome analyses, global histone methylation Western blots, and brain histology, respectively. We also thank Drs. Sally Camper, Stephen Parker, Stephanie Bielas, and Michael-Christopher Keogh, as well as the members of the Iwase and Bielas labs, for helpful discussions and critical review of the data. This work was funded by an NIH National Research Service Award T32-GM07544 (University of Michigan Predoctoral Genetics Training Program) from the National Institute of General Medicine Sciences (to CNV), an NIH National Research Service Award T32-HD079342 (University of Michigan Predoctoral Career Training in the Reproductive Sciences Program) from the National Institute of Child Health and Human Development (NICHD) (to CNV), University of Michigan Rackham Predoctoral Research Grants (to CNV), a Michigan Institute for Clinical and Health Research fellowship (Translational Research Education Certificate, supported by UL1TR000433 and UL1TR002240) (to CNV), a University of Michigan Rackham Predoctoral Fellowship award (to CNV), an Autism Science Foundation Predoctoral Fellowship award (to CNV), an NIH National Research Service Award. F31NS103377 from the National Institute of Neurological Disease & Stroke (NINDS) (to RSP), NIH NINDS Awards (R01NS089896 and R21NS104774) (to SI), Basil O’Connor Starter Scholar Research Awards from March of Dimes Foundation (to SI), and a Farrehi Family Foundation Grant (to SI).

## Author contributions

CNV, NCT, and SI conceived the study and designed the experiments. BR and KMC performed the mouse behavioral tests under the guidance of NCT. MCW oversaw dendritic morphometry analyses. CNV, KMB, PMG, and SI analyzed RNA-seq data. YAS performed global H3K4me3 analyses. RSP and SI performed H3K4me3 ChIP-seq and analyses. CNV performed the rest of experiments and analyses. YD and CEK provided key experimental recourse and made important intellectual contributions. CNV, MCW, RSP, CEK, NCT, and SI wrote and edited the manuscript.

---

**Supplementary Figure 1.**
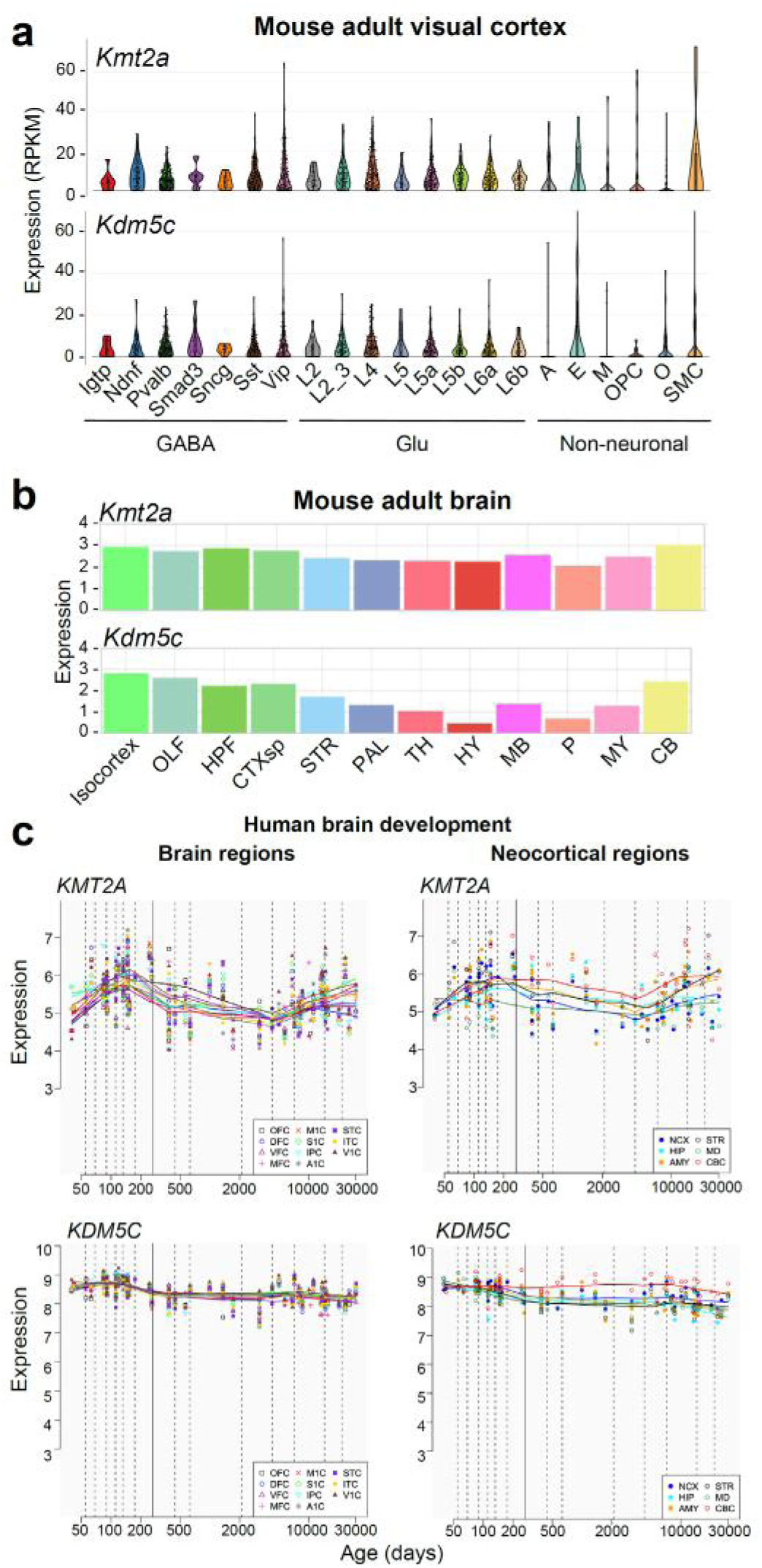
Expression of KMT2A and KDM5C. (A) Expression of *Kmt2a* and *Kdm5c*, from FACS-sorted single cells of mouse visual cortex, shown in reads per kilobase of transcript per million mapped reads (RPKM). Neuronal cells: GABAergic (GABA), Glutamatergic (Glu). Non-neuronal cells: astrocytes (A); endothelial cells (E); microglia (M), oligodendrocyte precursor cells (OPC); oligodendrocytes (O); smooth muscle cells (SMC). Image credit: Broad Institute “Single Cell Portal” transcriptome of adult mouse visual cortex (1). **(B)** Expression of *Kmt2a* and *Kdm5c* mRNA from adult mouse brain, shown in log2 of raw expression value from *in situ* hybridization. Brain regions: Isocortex, olfactory areas (OLF), hippocampal formation (HPF), cortical subplate (CTXsp), striatum (STR), pallidum (PAL), thalamus (TH), hypothalamus (HY), midbrain (MB), pons (P), medulla (MY), cerebellum (CB). Image credit: Allen Institute, Allen Mouse Brain Atlas (2004) (2). **(C)** Expression of *KMT2A* and *KDM5C* transcripts, from developing and adult human brains, shown in RPKM. Human development and adulthood were split into the following Periods: 1-7 fetal development; 8-9 birth and infancy; 10-11 childhood; 12 adolescence; and 13-15 adulthood. Image credit: Human Brain Transcriptome Atlas (3, 4)

**Supplementary Figure 2.**
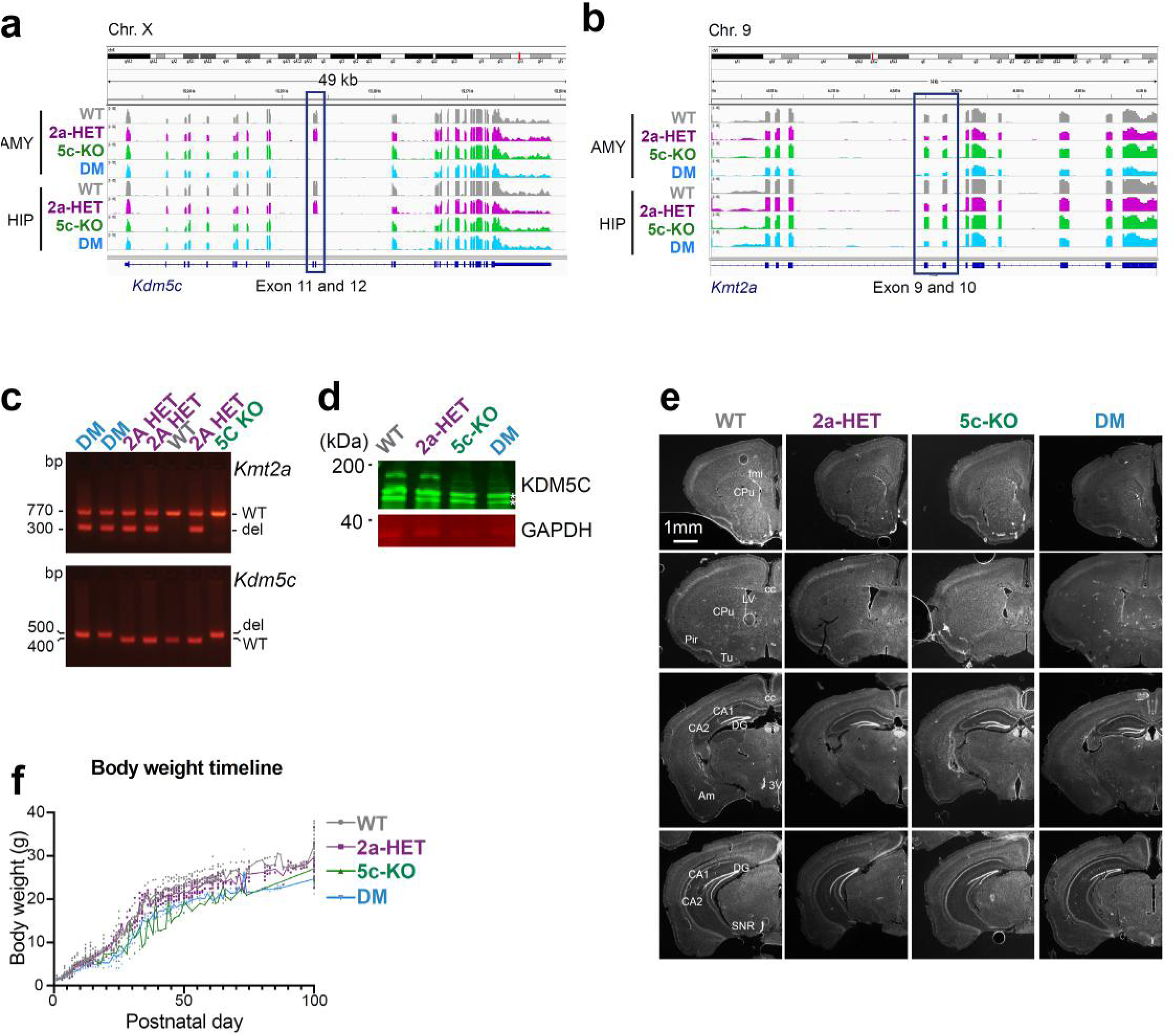
Basic features of mutant mice. **(A-B)** RNA-seq read coverage of *Kmt2a* **(A)** and *Kdm5c* **(B)** genes and targeted exons (highlight) confirmed the intended gene manipulations. **(C)** Genotyping using genomic DNA, confirming presence of *Kmt2a* and/or *Kdm5c* deleted alleles (“del”) only in appropriate genotypes. **(D)** Western blot for KDM5C protein. Stars indicate non-specific bands present in all samples. GAPDH shown for equal loading. **(E)** Serial brain sections 30 μm thick stained with DAPI to mark nuclei. Sections shown at Bregma regions 1.41, 0.49, −2.15, and −2.91 mm (top to bottom). Regions highlighted: anterior forceps of the corpus callosum (fmi), caudate putamen (CPu), corpus callosum (cc), lateral ventricle (LV), piriform cortex (Pir), olfactory tubercle (Tu), hippocampal fields CA1 and CA2, dentate gyrus (DG), anteromedial nucleus (AM), third ventricle (3V), substantia nigra pars reticularis (SNR). Scale bar: 1mm. **(F)** Body weight tracked from birth, postnatal day 1 (P1).

**Supplementary Figure 3.**
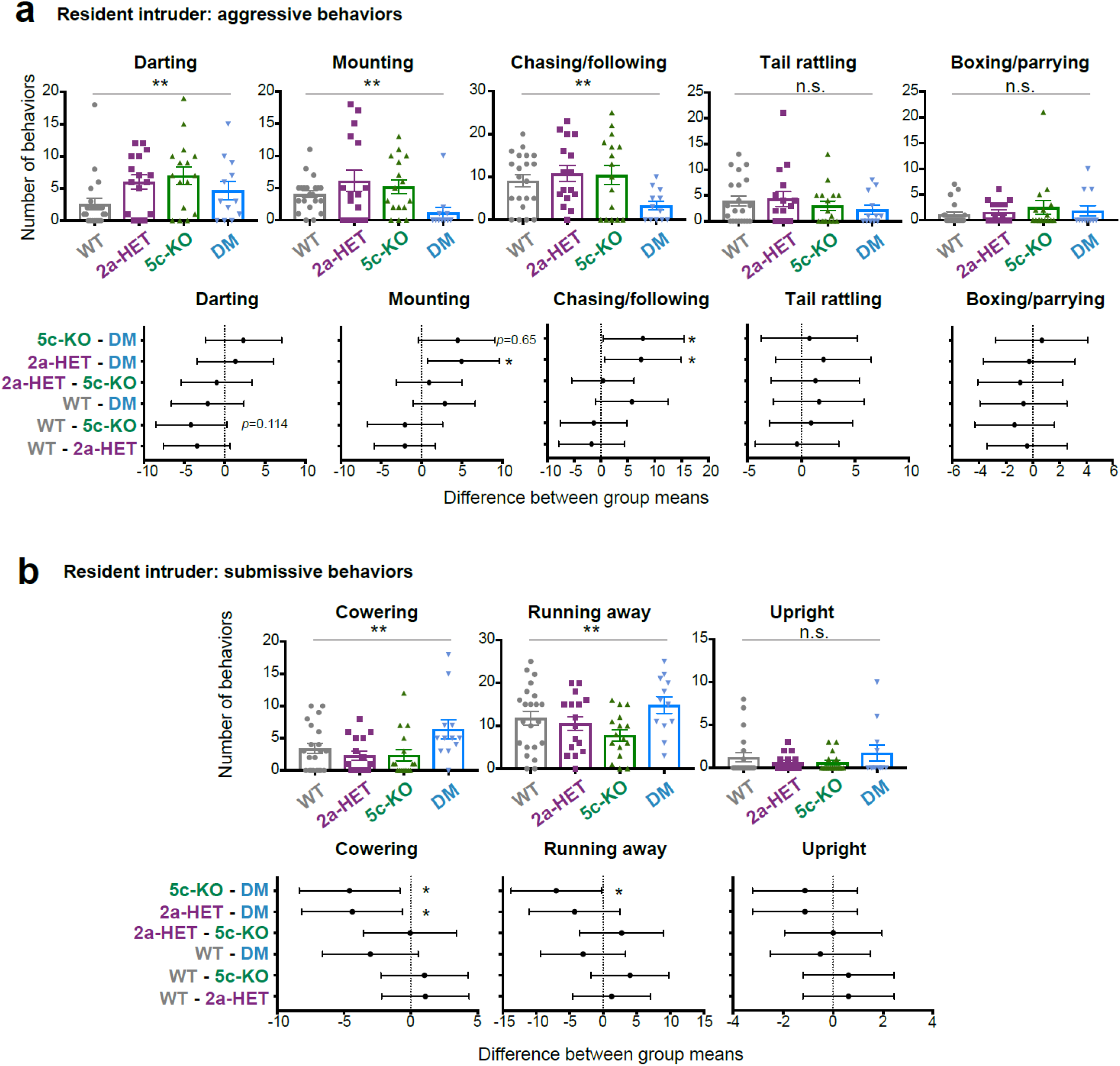
Individual behavior types in the resident intruder test. (A) Individual aggressive behaviors (mean ± SEM, \*\**p* < 0.01 in one-way ANOVA). N.S. depicts no statistical difference. Left panel: average number of all submissive behaviors (mean ± SEM, \**p* < 0.05 in one-way ANOVA). Right panel: Difference between group means of submissive behaviors (mean ± 95% confidence intervals, \**p*<0.05, \*\**p*<0.01). **(B)** Individual submissive behaviors (mean ± SEM, \*\**p* < 0.01 in one-way ANOVA). N.S. depicts no statistical difference. N=21 WT, N=16 *Kmt2a*-HET, N=16 *Kdm5c*-KO, and N=12 DM animals were used for all studies. Differences between group means all aggressive **(A)** and submissive **(B)** behaviors (mean ± 95% confidence intervals, \**p*<0.05).

**Supplementary Figure 4.**
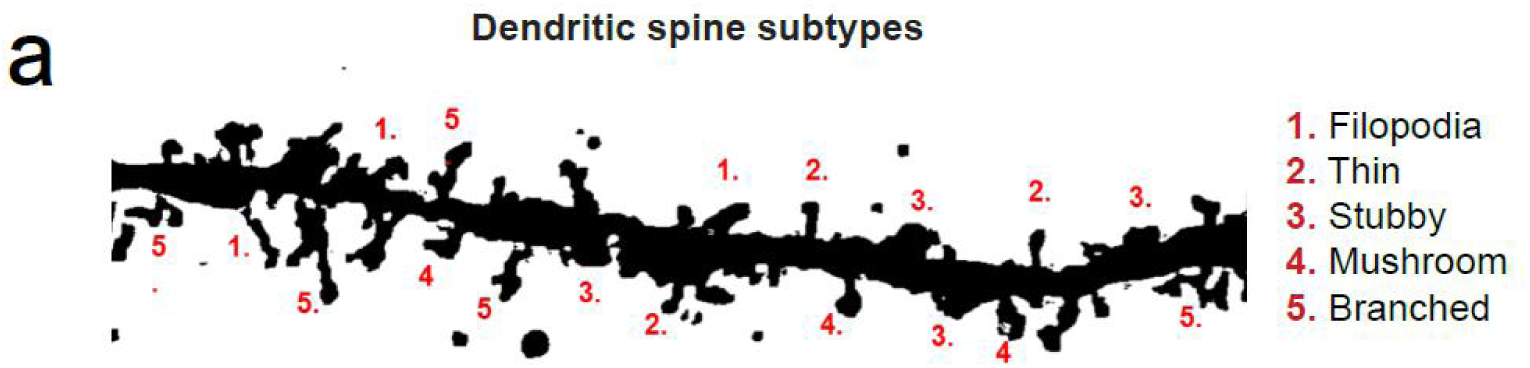
Schematic of dendrite spine subtype analysis. (A) Projection image of a dendritic segment from a series of Z stack images derived from BLA pyramidal cells. Numerical marks adjacent to corresponding spine subtypes, represented as: 1. Filopodia, 2. Thin, 3, Stubby, 4. Mushroom, and 5. Branched.

**Supplementary Figure 5.**
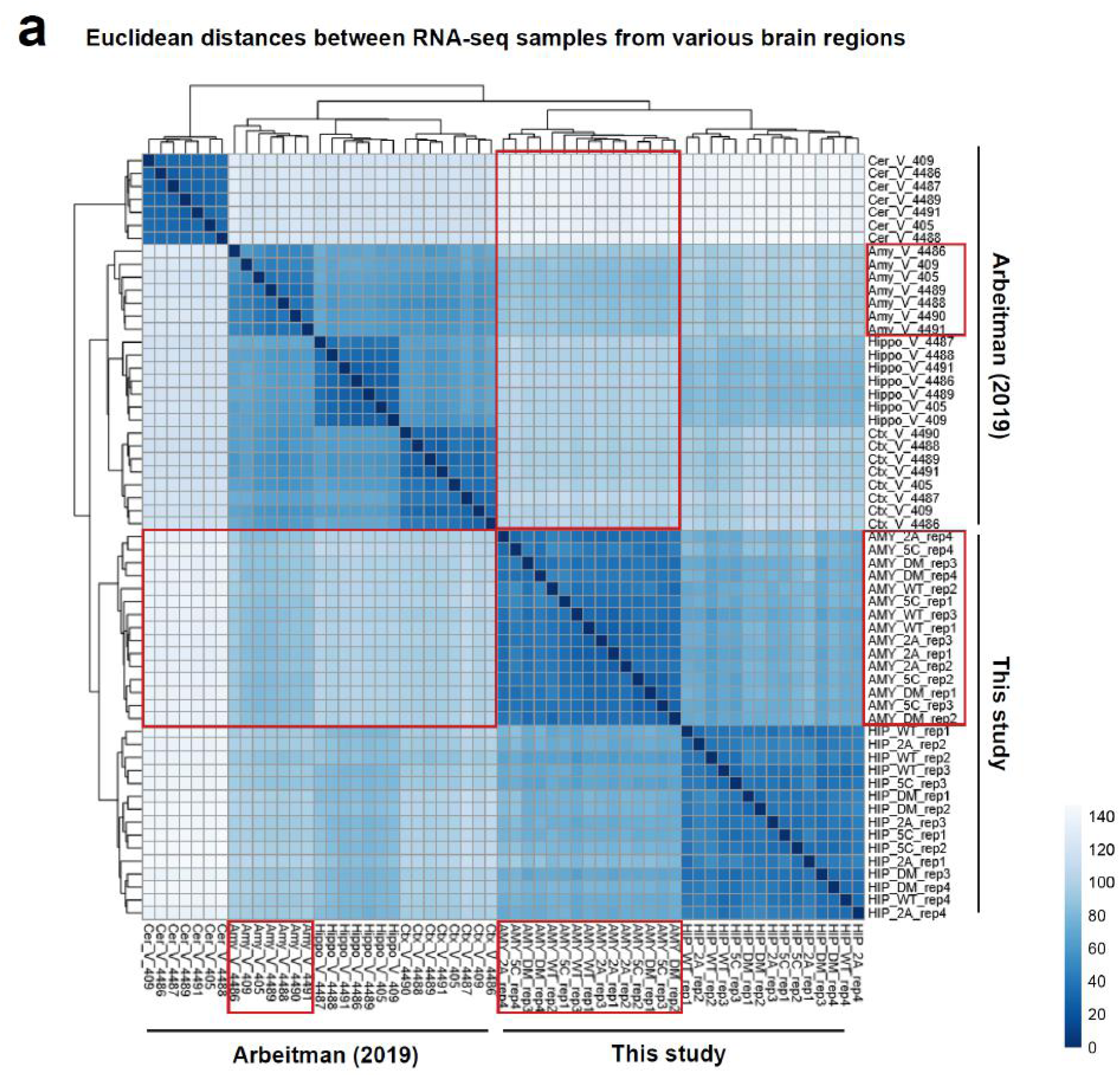
Validation of brain microdissection. (A) To validate accuracy of our brain microdissection, we compared our RNAseq data of the hippocampus (HIP) and amygdala (AMY) with those of the Arbeitman paper (5), which involved the hippocampus (Hippo), amygdala (Amy), cerebellum (Cer), cerebral cortex (Ctx). Euclidian distance between all combination of RNA-seq data are plotted (see method). Samples from this study and the Arbeitman study are clustered separately, likely due to difference in experimental procedures and/or sex of mice. Our study used adult male mice, while the Arbeitman study used adult females. Nonetheless, our amygdala samples showed the shortest distance to the Arbeitman amygdala data compared to any other brain tissues (red rectangles). Likewise, our hippocampus data are closest to the Arbeitman hippocampus data among the four brain regions. The data demonstrate that the microdissection of brain regions in the two studies are consistent with each other.

**Supplementary Figure 6.**
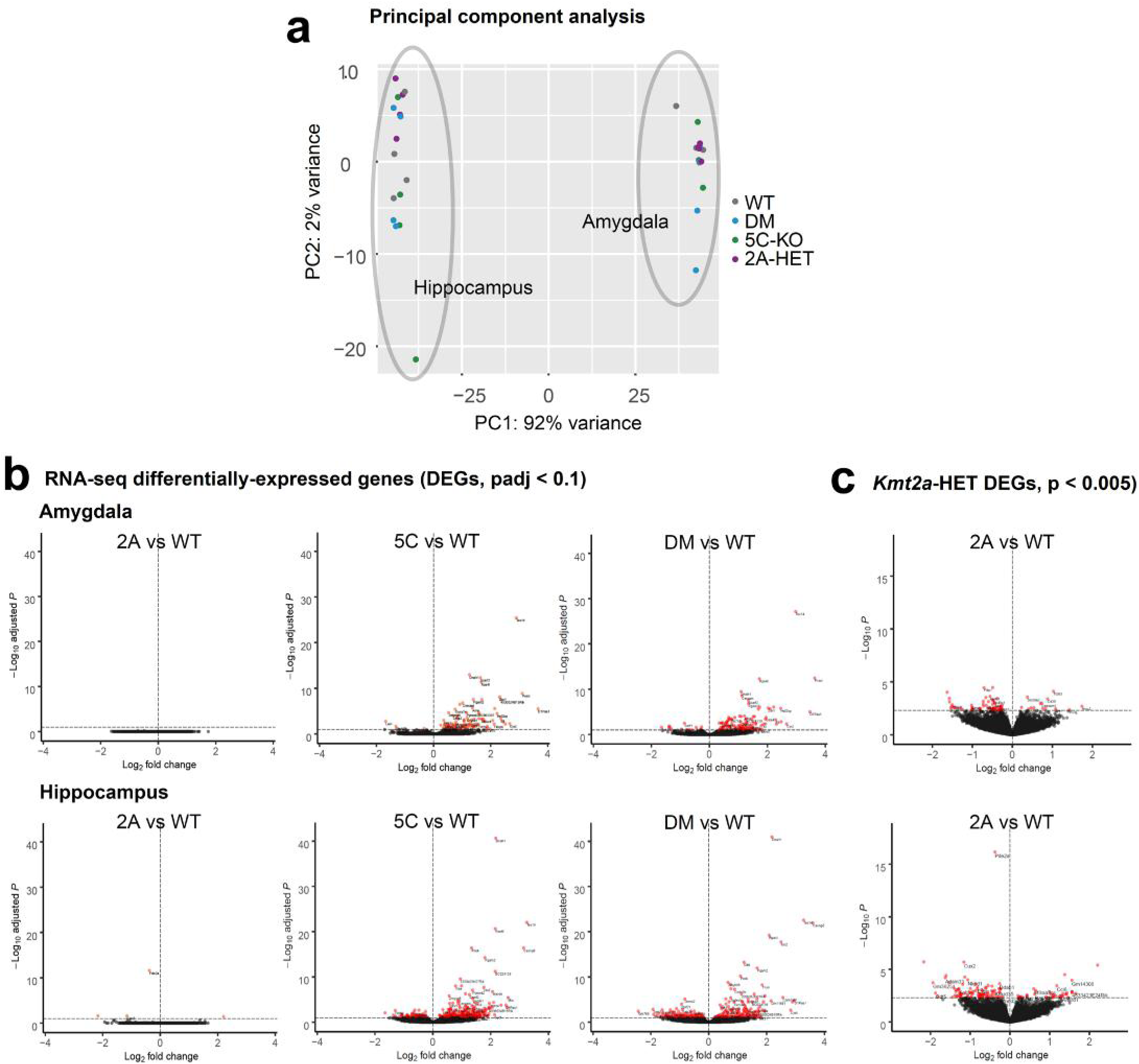
Basic analysis of RNA-seq data. (A) PCA analysis of RNA-seq libraries. Tissue types are a stronger segregating factor than genotypes. **(B)** Volcano plot representation of differential expressed genes (DEGs) in mutant vs WT comparisons. Red dots: DEGs with *padj* < 0.1 (see Table S1). **(C)** Mildly dysregulated genes in Kmt2a-HET were recovered with a relaxed threshold.

**Supplementary Figure 7.**
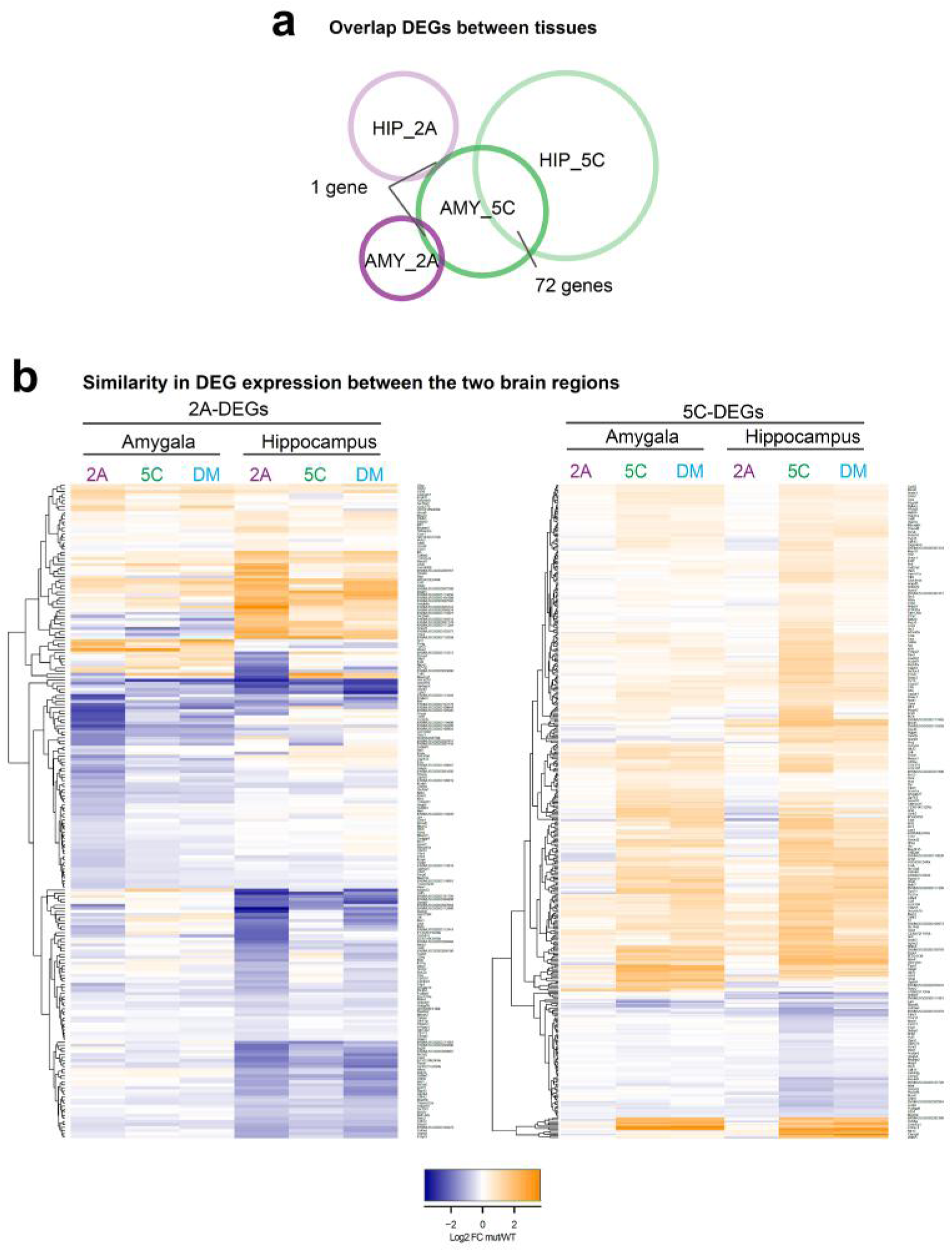
Similarity of gene misregulation between the amygdala and hippocampus. (A) Overlap of single mutant DEGs between the amygdala (AMY) and hippocampus (HIP). 5C-DEGs overlap between the two brain regions. **(B)** Heatmap representation of log2 fold changes of 2A-DEGs (left) and 5C-DEGs (right). Overall the patterns of gene misregulation are similar in the two brain regions.

**Supplementary Figure 8.**
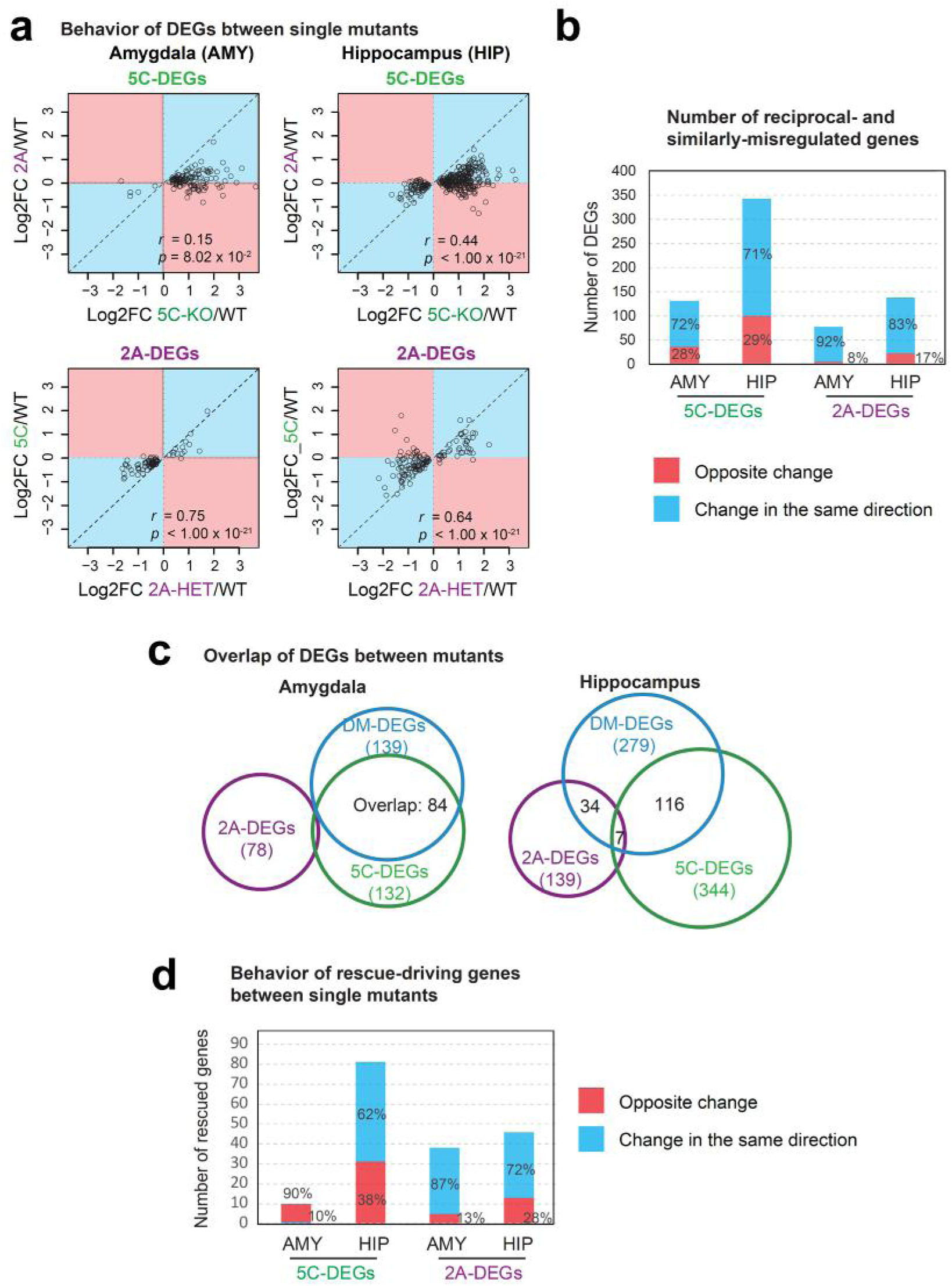
Direction of gene misregulation between *Kmt2a*-HET and *Kdm5c*-KO and their overlap in DM brain tissues. (A) Behavior of single-mutant DEGs in the other single mutant. Log2 fold changes of DEGs found in a single mutant were plotted as a function of log2 fold changes in the other single mutant. Red shade covers genes that are regulated in the opposite direction between *Kmt2a*-HET and *Kdm5c*-KO brain tissues. Blue shade covers deregulated genes in the same direction. **(B)** The larger number of genes is dysregulated in the same direction between the two single mutants. **(C)** Overlap of DEGs between mutants. 5C- and DM-DEGs overlap substantially.

**Supplementary Figure 9.**
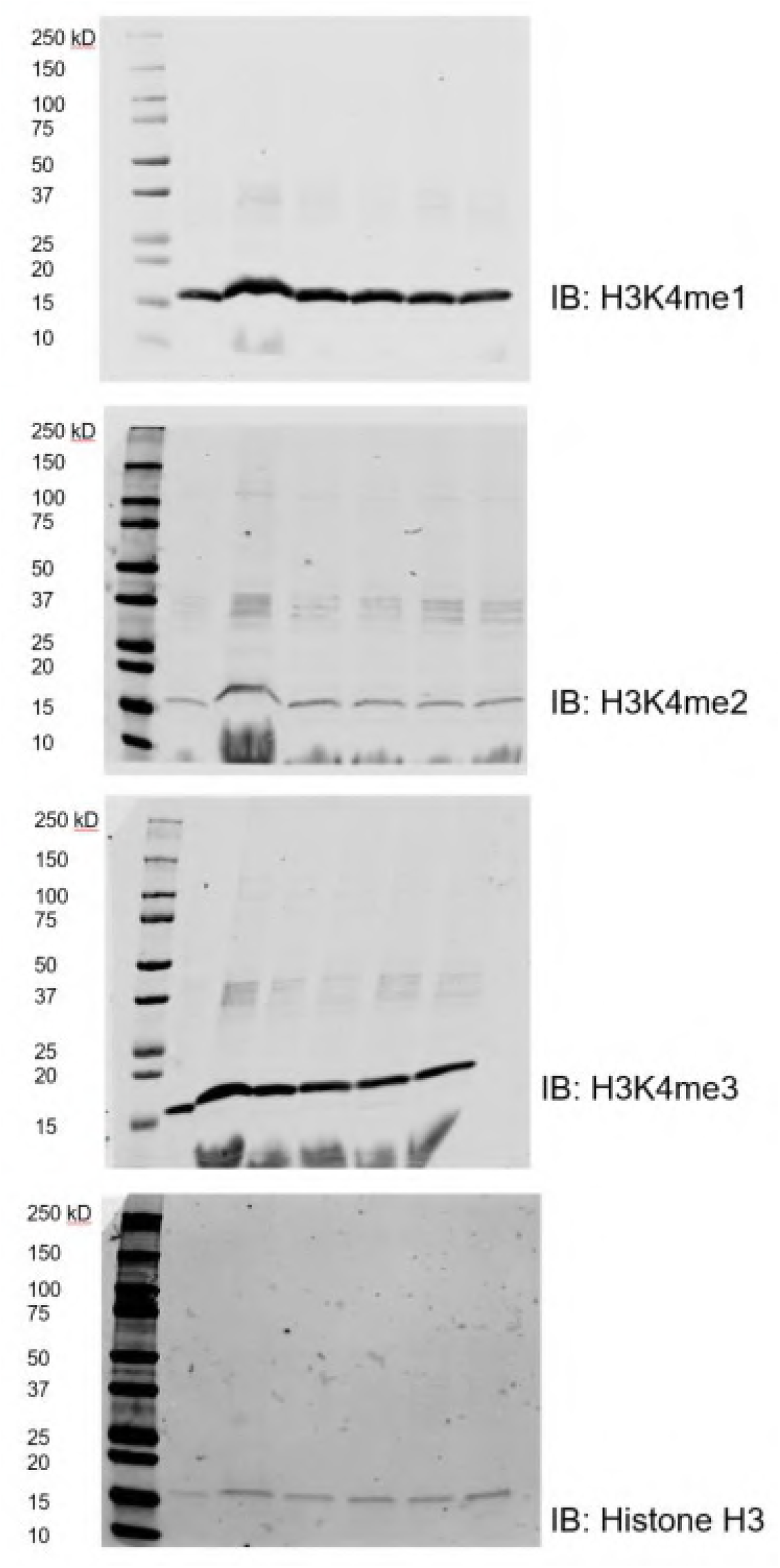
Full blots of immunoblot (IB) analyses shown in Figure 6a.

**Supplementary Figure 10.**
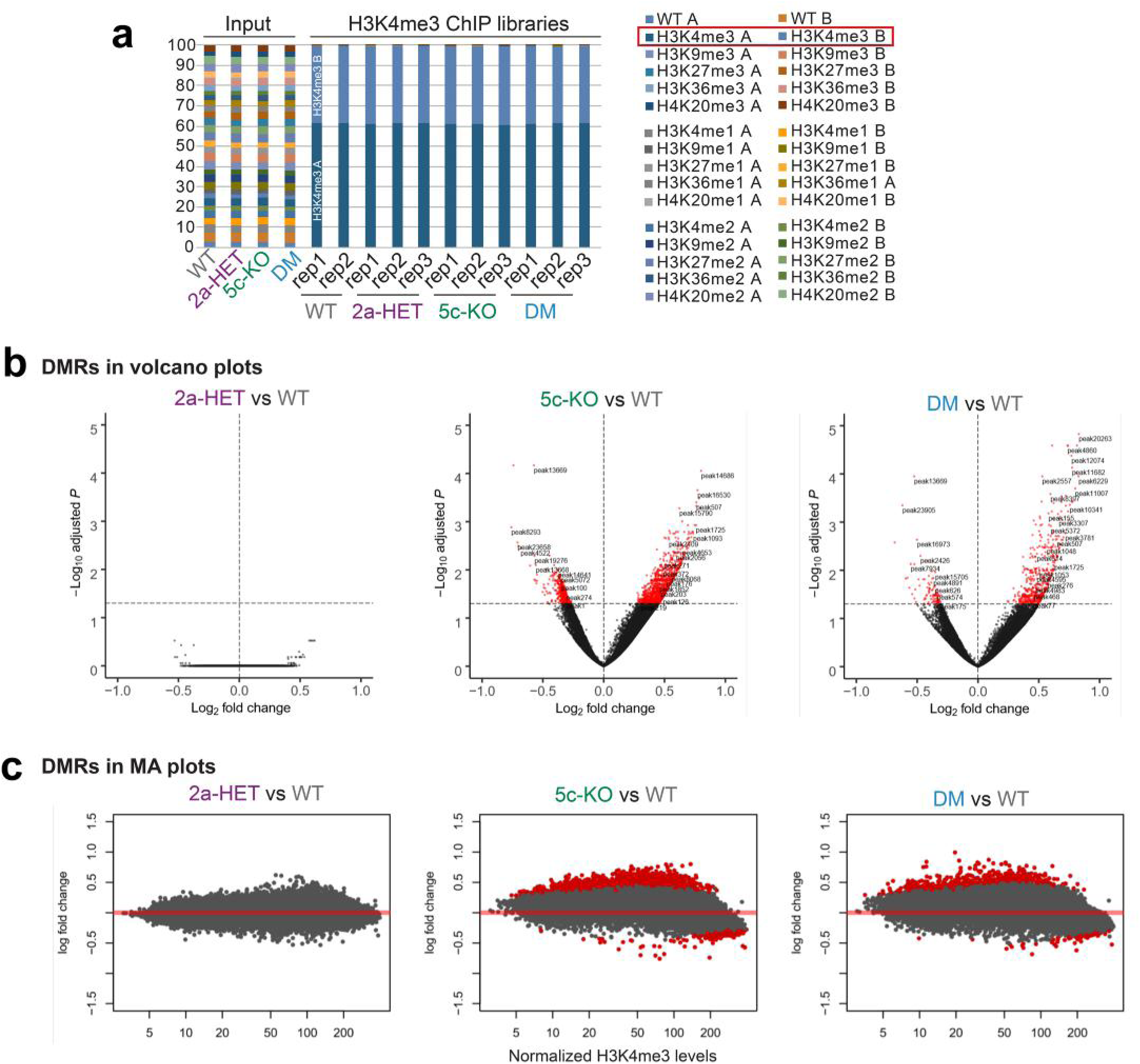
Basic characterization of H3K4me3 ChIP-seq data. (A) Validation of H3K4me3 ChIP-seq specificity. Barcode reads originating from spike-in nucleosomes were counted. The two synthetic nucleosomes with the H3K4me3 barcodes dominated all ChIP samples, with H3K4me1/2 nucleosomes rarely detected. **(B)** Volcano plot represents the statistical significance of differentially methylated regions (DMRs). **(C)** MA plots of DMRs revealed signal intensity-dependent misregulation of H3K4me3 in *Kdm5c*-KO, where hypomethylated regions in the mutant were highly methylated in WT.

**Supplementary Figure 11.**
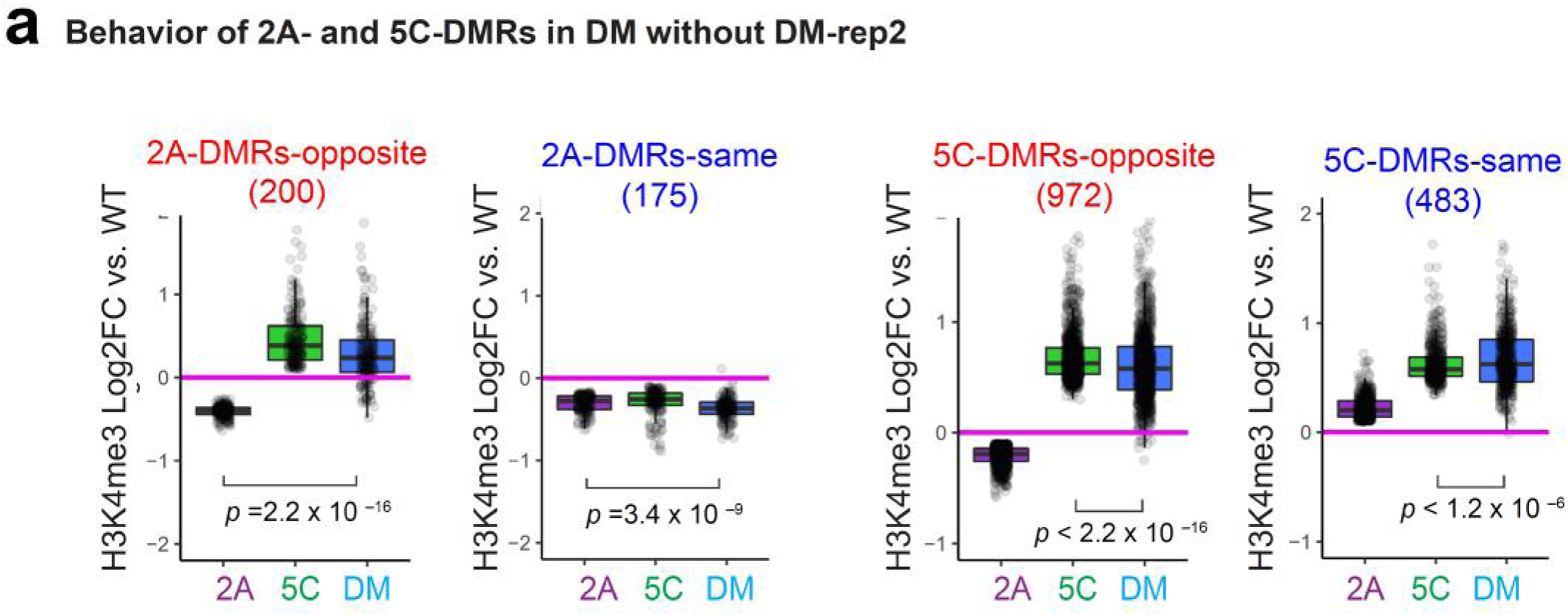
Rescue effect without DM rep2. (A) DM rep2 showed strong rescue effect (Figure 6B). To test if H3K4me3 misregulations were alleviated in other DM replicates, we removed DM rep2 and then examined the behavior of single-mutant DMRs in DM. Log2 fold change of DMRs relative to WT were plotted across the three mutants. Boxplot features: box, interquartile range (IQR); bold line, median; gray dots, individual genes. Associated *p* values result from Wilcoxon signed-rank tests. Rescue effects were still evident in DM and also dependent on the direction of misregulation between the single mutants.

**Supplementary Figure 12.**
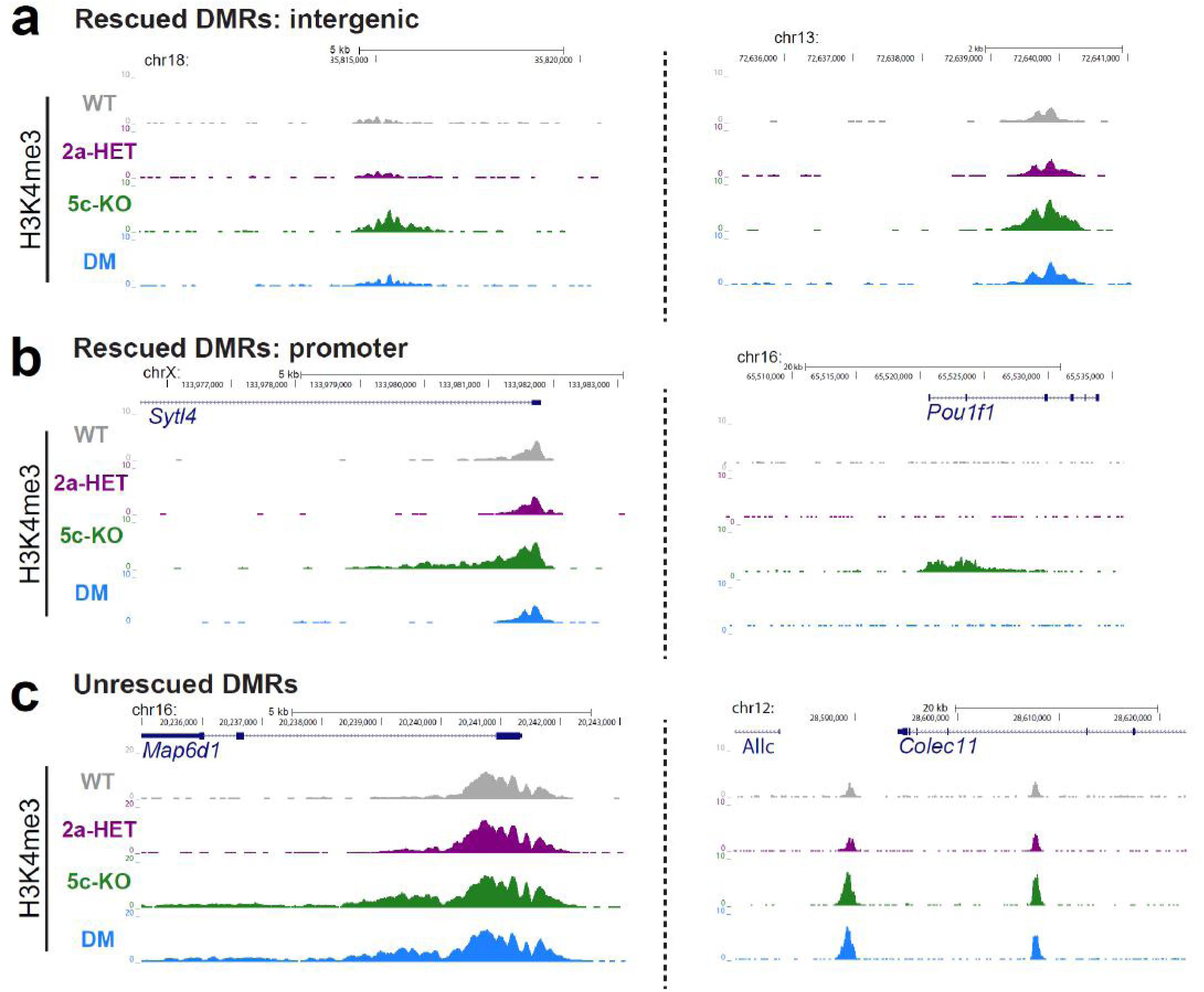
Representative loci found in the H3K4me3 ChIP-seq analysis. (A) Representative genome browser view of two representative loci for each of the major genome areas: rescued intergenic DMRs **(A)**, rescued promoter DMRs **(B)**, un-rescued DMRs **(C)**. Represented H3K4me3 patterns are averaged signals of replicates that are normalized to read depth and spike-in nucleosome signals.

